# Maternal histone methyltransferases antagonistically regulate monoallelic expression in *C. elegans*

**DOI:** 10.1101/2024.01.22.576748

**Authors:** Bryan Sands, Soo R. Yun, Junko Oshima, Alexander R. Mendenhall

## Abstract

Undefined epigenetic programs act to probabilistically silence individual autosomal alleles, generating unique individuals, even from genetic clones. This sort of random monoallelic expression can explain variation in traits and diseases that differences in genes and environments cannot. Here, we developed the nematode *Caenorhabditis elegans* to study monoallelic expression in whole tissues, and defined a developmental genetic regulation pathway. We found maternal H3K9 histone methyltransferase (HMT) SET-25/SUV39/G9a works with HPL-2/HP1 and LIN-61/L3MBTL2 to randomly silence alleles in the intestinal progenitor E-cell of 8-cell embryos to cause monoallelic expression. SET-25 was antagonized by another maternal H3K9 HMT, MET-2/SETDB1, which works with LIN-65/ATF7ZIP and ARLE-14/ARL14EP to prevent monoallelic expression. The HMT-catalytic SET domains of both MET-2 and SET-25 were required for regulating monoallelic expression. Our data support a model wherein SET-25 and MET-2 regulate histones during development to generate patterns of somatic monoallelic expression that are persistent but not heritable.

## Introduction

When accounting for variation in traits, scientists bin variation into genes and environments. But there is a third component that accounts for biological variation that is considered nongenetic and nonenvironmental^1^. In model animal research, it is possible to hold genes and environments as constants, so the remaining variation in traits accounts for this nongenetic, nonenvironmental variation, which can be substantial^2^. Terms like incomplete penetrance and variable expressivity were coined to explain the observed variation in mutant phenotypes in isogenic populations of flies in homogeneous environments^3–5^. One of the major ways this intrinsic variation in discrete or complex traits manifests is through intrinsic variation in gene expression^6,7^. One major cause of intrinsic variation in gene expression is through endogenous silencing^8–10^. In humans, the endogenous silencing of genes works through random silencing of individual autosomal alleles early in development, evidenced by experimental studies and non-Mendelian escape from genetic disease, discussed further below. This kind of intrinsic variation in gene expression can account for incomplete penetrance, missing heritability, and nongenetic, nonenvironmental variation in complex traits, including the manifestation of chronic diseases. We refer to this kind of probabilistic silencing of alleles as monoallelic expression (MAE), reviewed in^8–10^.

### Monoallelic expression in humans

Random autosomal MAE is consequential, but not entirely understood^8–10^. Other forms of allelic silencing are associated with biological processes including imprinting, the selection of alleles of odorant receptors, and the expression of antibodies^9^. The probabilistic silencing that occurs on autosomal alleles was found to be widespread^11^ and, since it is propagated through tissues, was detectable with both bulk RNA-seq^12,13^ and ChIP-seq^14,15^. Recent analysis of GTEX RNA-seq data has shown that genes with MAE potential are overwhelmingly enriched for effects on aging and chronic disease like cancer^13^. Moreover, MAE operates on a continuum, from fully monoallelic to biallelic, and the silencing of alleles seems to be persistent, but not heritable between generations^9,16^.

### Monoallelic expression in human genetic disease and cancer

There are several cases of monoallelic expression affecting the penetrance of genetic disease, cited in these reviews^8–10,17^. One example is a case study of a family that harbored a dominant-negative mutation in *PIT1*^18^. In the three heterozygous individuals that harbored the dominant-negative disease allele (grandmother, father, and daughter), only the daughter was affected by disease. Analysis of RNA showed that the pathological dominant-negative disease allele was silenced in the disease-free family members (grandmother and father), and expressed in the affected daughter.

The variable onset of genetic diseases can also be explained by MAE. Genetic diseases like *ACTG2* related visceral myopathy have variable onset, with some people remaining symptom-free until later in life^19^. This late onset is consistent with the idea that desilencing of pathological alleles could cause the later onset disease. Moreover, the silencing of non-pathological alleles in people that are heterozygous for recessive pathological alleles can lead to manifestation of recessive disease that Mendelian genetics would not predict. Finally, in addition to changes in biochemical activity conferred by alleles with different sequence, silencing of alleles will generally lower protein dosage/activity (some genes have compensation mechanisms^20^).

Monoallelic expression of several different genes in tumors can act as prognostics in cancer medicine^21–28^. Monoallelic expression can mean increased life expectancy or decreased life expectancy, depending on the particular scenario. In some cases, it may be that monoallelic expression simply reduces the dosage of a metabolically advantageous protein, like IDH1^21^. In other cases, monoallelic expression might mean an increased likelihood of immune escape. Homologs for genes we identified as regulating MAE in this report affect the immunogenicity of tumors^29^ and cancer cell lines^30^.

### *C. elegans* as a model to study random autosomal monoallelic expression

We have developed *C. elegans* as a model to study monoallelic expression *in vivo*. This model utilizes two distinctly colored fluorescent alleles integrated at the same locus to visualize and quantify differences in allele expression at the protein level in individual somatic cells. The system addresses a gap between sequencing based approaches to study monoallelic expression and medical cases of monoallelic expression^31^. Moreover, there are technical advantages to using a nematode model to study MAE. These include a transparent soma, the ease of conducting genetic screens, a short life cycle, and a fixed mitotic lineage. Together, these properties allow us to directly visualization the expression of individual alleles in multiple tissues, assess heritability, distinguish between steady-state expression and transcriptional bursting, and to infer developmental events from somatic expression patterns^32–39^.

Here, we performed a targeted screen for negative regulators of MAE and identified the H3K9-specific histone methyltransferase (HMT) *met-2/SETDB1*^40^ as a major negative regulator. Further analysis of MET-2 associated genes revealed that another H3K9-specific HMT, SET-25^40–42^, antagonistically regulated MAE with MET-2, similar to their genetic effects on germline immortality^43^. Loss of MET-2, or just its catalytic HMT SET domain, promoted MAE, while loss of SET-25 activity, or just its SET domain, prevented MAE, resulting in biallelic expression (BAE). *met-2* mutant animals showed a pattern of monoallelic expression that was persistent throughout adulthood, but not heritable. Additionally, animals lacking functional catalytic domains of MET-2, SET-25, or both, were able to use RNA interference to silence somatic genes, demonstrating that the genes regulating MAE are not required for regulating RNA interference. Reciprocal cross experiments showed that maternal MET-2 and SET-25 act in the E-cell of the 8-cell embryo to regulate MAE. Our results support a model wherein MET-2 and SET-25 competitively regulate histone marks to control nucleosome occupancy and silencing of alleles of genes subject to monoallelic expression.

## RESULTS

### Targeted screening identifies MET-2/SETDB1 as a negative regulator of **monoallelic expression**

We have previously developed a reporter allele system to quantify expression of alleles at the protein level in single cells of whole tissues in *C. elegans*^32^. Our system utilizes two differently colored fluorescent alleles, such as *hsp-90* alleles, that are expressed from the same locus in the somatic cells of *C. elegans*^32^. The animals are propagated by picking self-fertile hermaphrodites that are heterozygous for the reporter alleles at each generation. Under standard conditions (OP50-seeded NGM at 20°) intestine cells express *hsp-90* reporter alleles in a biallelic fashion, with occasional instances of allele bias or monoallelic expression detected^32^.

Here, we conducted a targeted reverse genetic screen to identify negative regulators of MAE. To do this, we subjected worms that are heterozygous for *hsp-90* reporter alleles to two generations of RNAi (Figure 1a), loaded them into microfluidics devices, and imaged them by confocal microscopy (Figure 1b). Worms expressing *hsp-90* reporter alleles on empty vector (EV) were used as a control. We screened chromatin-modifying genes^44^, including SET-domain proteins, DNA modifiers, known or predicted histone modifiers, and proteins associated with the endogenous silencing machinery, for increases in MAE (Table 1).

**Figure 1.**
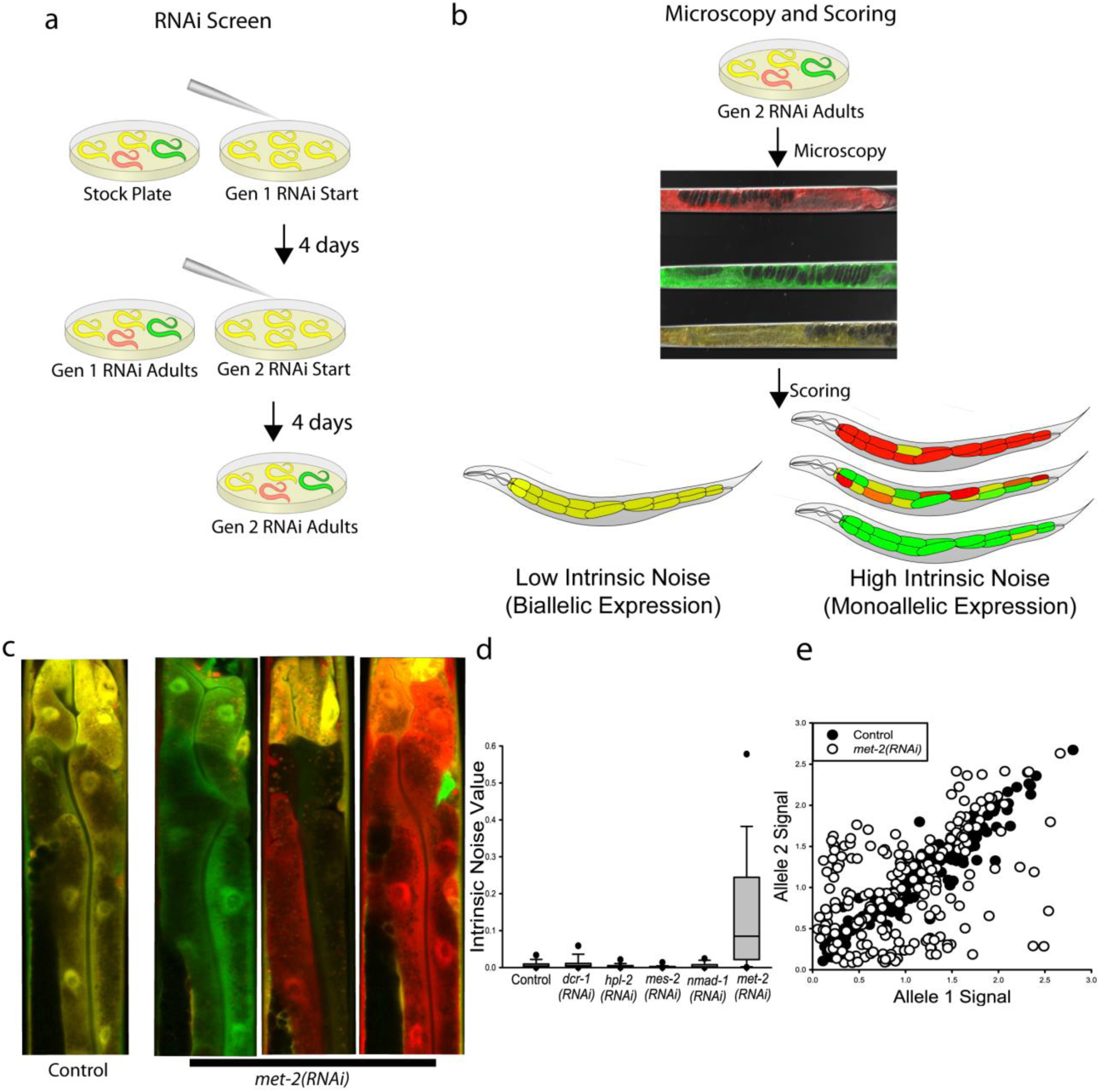
MET-2 negatively regulates monoallelic expression. **a)** shows a schematic overview of how we implemented RNAi in animals that were heterozygous for reporter alleles for two generations. **b)** shows how we mounted animals for confocal microscopy and cartoons of the spectrum of phenotypes we would observe. **c)** shows actual merged confocal images of the “torso” of control animals and *met-2(RNAi)* animals. The torso image is the section of the animal that fits into the field of view of our 40X objective and contains cells in the first three rings of the intestine. **d)** shows boxplots of intrinsic noise, a quantification of allele bias/MAE, calculated from the intestine cell allele expression values for control animals, a few of the genes we screened that did not alter MAE, and *met-2*. **e)** shows intestine cells plotted by allele expression level from control and met-2(RNAi) animals. There was a significant increase in intrinsic noise for met-2 animals, compared to control animals (0.00371); *P* < 0.05; Kruskal-Wallis One Way Analysis of Variance on Ranks followed by Dunn’s Procedure, over 200 cells per group, three independent experiments. See also Supplementary Information for additional statistical details and Supplemental Figures 1&2, showing developmental events indicated by monoallelic expression patterns and detailing additional RNAi test results.

**Table 1:**
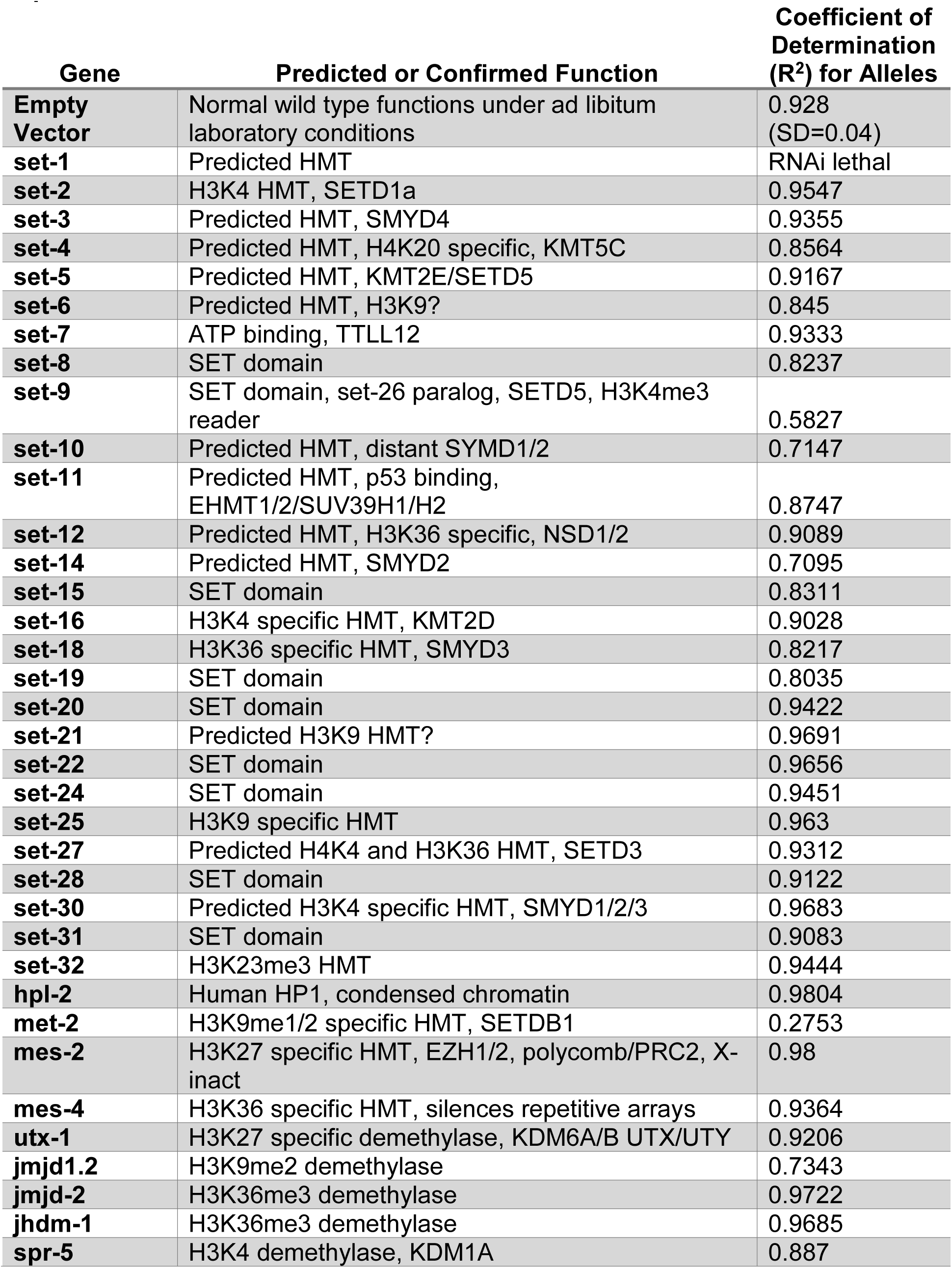
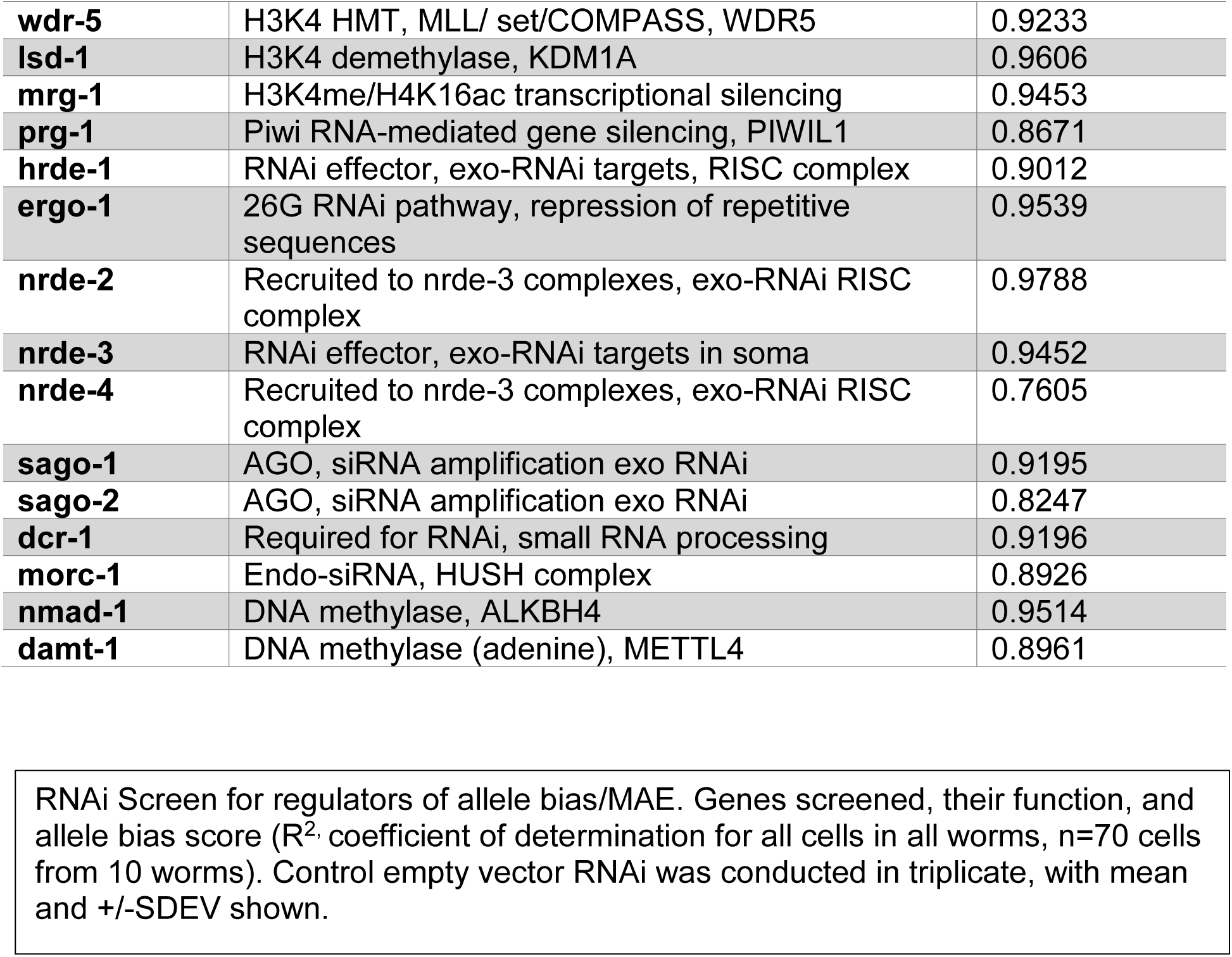
Targeted Genetic Screen for Negative Regulators of Monoallelic Expression.

Because *hsp-90* reporter allele expression usually results in a biallelic expression pattern, we screened for animals that showed extreme MAE. In our screen, only *met-2* showed extreme MAE, with *met-2(RNAi)* animals showing a pattern of MAE across most somatic cells, including those that we quantified in the intestine (Figure 1c-e). *met-2* encodes a conserved H3K9 HMT that is homologous to human *SETDB1*^40–42^ and is generally associated with silencing, not preventing it. Figure 1d shows the intrinsic noise values for control and *met-2(RNAi)* compared to a subset of other genes from the screen. To validate the initial *met-2(RNAi)* screen result, we conducted additional RNAi experiments and found loss of MET-2 activity resulted in a large, significant increase in monoallelic expression, quantified as intrinsic noise (Figure 1e and Supplemental Figure 1). Scatter plots of individual intestine cells plotted by allele expression values from control and *met-2(RNAi)* animals are shown in Figure 1e. These RNAi experiments confirmed that there is a significant increase in MAE when MET-2 activity is reduced via RNAi.

To determine if a null mutant would recapitulate the RNAi phenotype, we created a new knockout allele of *met-2* using CRISPR/Cas9 to insert multiple stop codons into the first exon of the *met-2* reading frame (see Methods and Supplemental Table 3). The *met-2(wam007)* animals phenocopied the *met-2(RNAi),* resulting in significantly higher median intrinsic noise, or, significantly increased MAE, compared to the control animals (Figure 2a).

**Figure 2.**
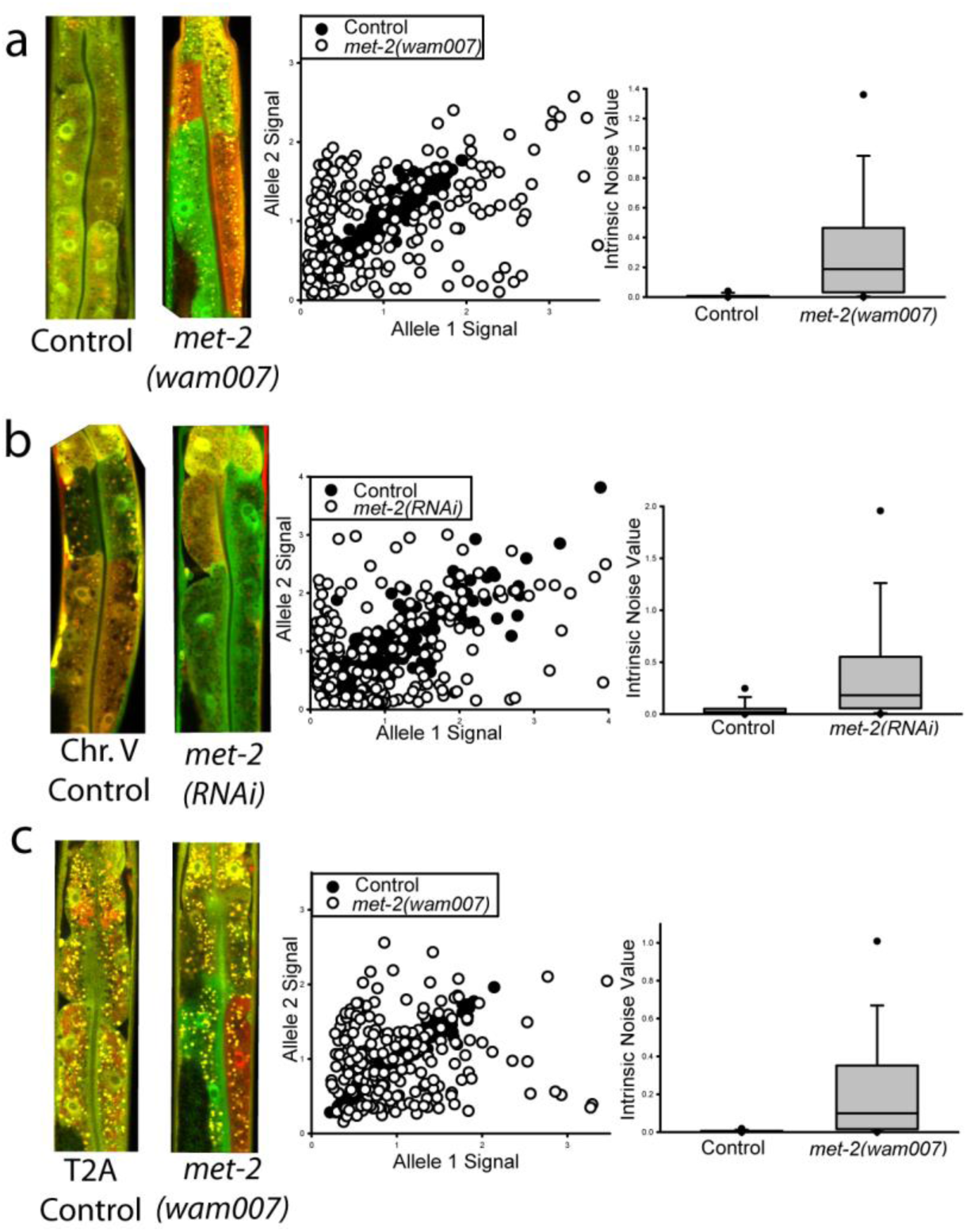
MET-2 regulates monoallelic expression when coding sequence and locus are changed. Each row, **a-c**, shows: 1) images of control and experimental animals’ “torso”, comprised of cells forming the first three “rings” of the *C. elegans* intestine; 2) scatter plots of control and experimental group intestine cells plotted by allele expression values from all three independent experiments; 3) boxplots of control and experimental cell intrinsic noise values, a measure of how monoallelic or biallelic expression is. **a)** shows that *met-2(wam007)* null mutant animals’ intestine cells had a significantly higher median intrinsic noise of 0.187 compared to 0.00232 for control animals’ cells; *P* < 0.001, Mann-Whitney Rank Sum Test, over 200 cells from each group, three independent experiments. **b)** shows that *met-2* regulates MAE for the same gene, but at a different noisier locus on Chromosome V. The left panel shows images of animals expressing the *hsp-90* reporter alleles from Chromosome V. The middle panel shows the scatter plots of cells from control animals and *met-2(RNAi)* animals. The right panel shows that there was a significantly higher intrinsic noise for alleles in the met-2(RNAi animals compared to the control animals; *met-2(RNAi)* 0.180 vs control 0.0196, *P* < 0.001, Mann-Whitney Rank Sum Test, three independent experiments. **c)** shows that *met-2* regulates MAE for *hsp-90* reporter alleles that also contain the full coding sequence (introns and exons) for *hsp-90.* Boxplots show that the intestine cells of *met-2(wam007)* animals had significantly higher intrinsic noise for full length coding sequence *hsp-90* reporter alleles than control animals; 0.100 median intrinsic noise for *met-2(wam007)* animals’ cells vs 0.00163 for control animals, *P* < 0.001, Mann-Whitney Rank Sum Test, over 200 cells from each group, three independent experiments.

Previously, we found that locus had an effect on monoallelic expression of reporter alleles. In that study, we moved reporter alleles from a locus on Ch II to an expression-permissive locus on Ch V, and found that, despite expression levels being similar, MAE was significantly increased^32^. Therefore, we tested the effect of *met-2* on the same *hsp-90* reporter alleles at the noisier locus on Chr. V^32^. Similar to what we found with reporter alleles on Ch II, *met-2(RNAi)* animals expressing the reporter alleles from Chr. V had a significant increase in MAE compared to EV controls (Figure 2b). These results show that *met-2* negatively regulates MAE in two different chromatin contexts with different baseline MAE potential^32^.

In a prior report, we found changes to coding sequences did not affect MAE^32^, so we hypothesized that *hsp-90* reporter alleles with additional coding sequence would still respond to *met-2*. We changed the coding sequence from fluorescent proteins to the full length *hsp-90* coding sequence, including the native promoter and introns. We tagged the full length coding sequence with a *T2A* peptide followed by either mEGFP or mCherry, and inserted the reporter alleles at the Chr. II locus^32^. We then used CRISPR/Cas9 editing to generate the *met-2* knockout allele *met-2(wam007)* in these animals and measured intrinsic noise compared to unedited controls. In worms with full length *hsp-90* alleles and *met-2(wam007),* intrinsic noise was significantly higher than in controls, shown in Figure 2c. Taken together, we identified *met-2* as a negative regulator of MAE of *hsp-90* reporter alleles, whether the reporters were inserted at different loci or had different coding sequences (see Materials and Methods for details).

### MET-2 regulates MAE in a gene specific fashion

Monoallelic expression of different genes may be caused by different cis elements in a gene, and can be regulated in many ways. In a previous investigation, we found that MAE was regulated by promoter and not influenced by coding sequence^32^, consistent with bioinformatic findings^45^. Other studies have found that different monoallelically expressed genes can have different modes of epigenetic regulation^12,14,15,20,46^. To test the hypothesis that MET-2 regulates MAE in a gene or promoter specific fashion, we built additional reporter alleles and tested them for response to *met-2* RNAi or *met-2* knockout.

First, we examined the *vit-2* promoter, which drives extremely high expression of yolk protein during egg production^47,48^. We used CRISPR/Cas9 to generate *met-2(wam007)* in a *vit-2* reporter allele strain. We found no significant difference in median intrinsic noise for *vit-2* reporter alleles between *met-2(wam007)* and control worms, shown in Figure 3a.

**Figure 3.**
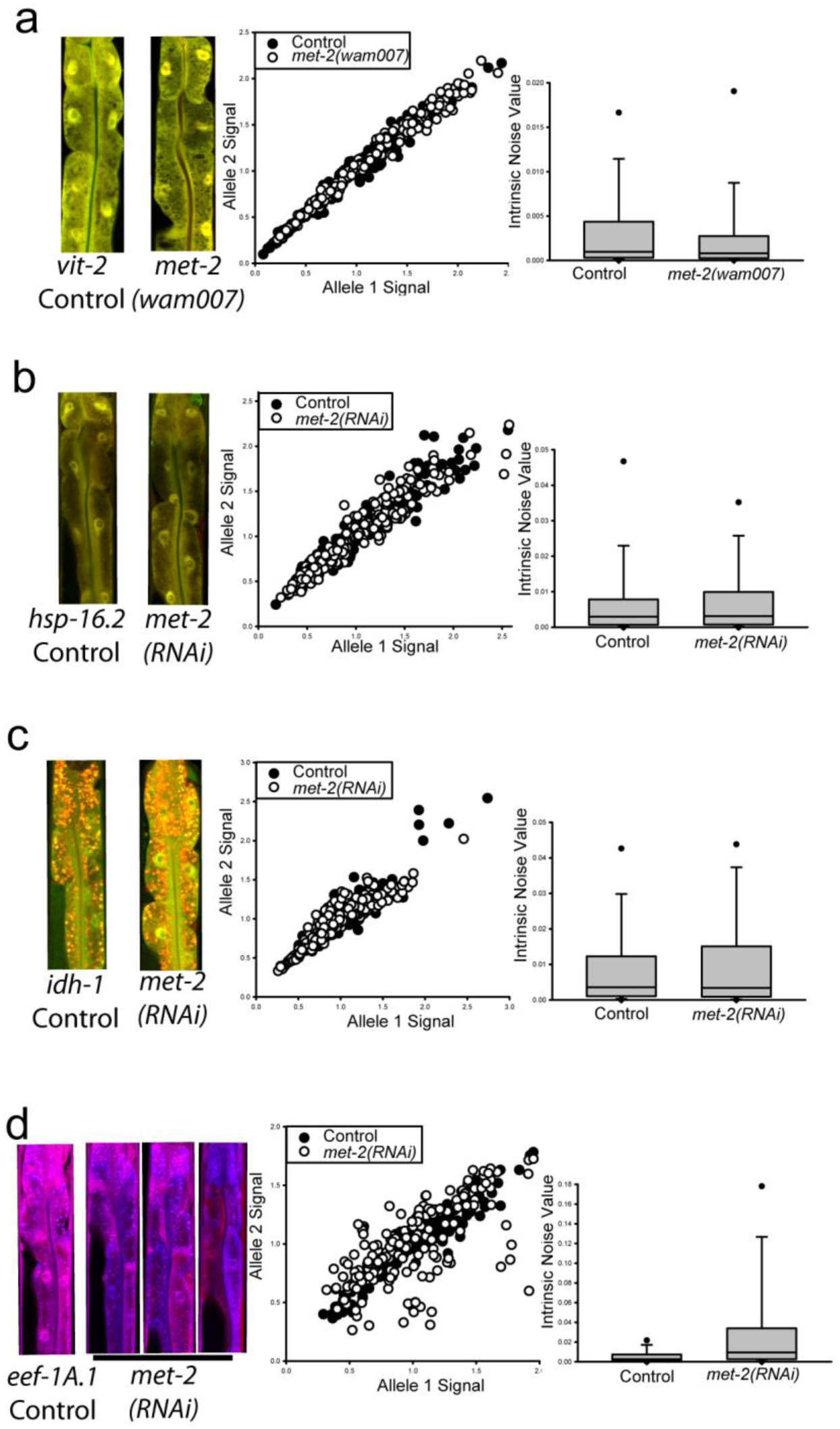
MET-2 negatively regulates monoallelic expression in a promoter/gene specific fashion. Images, scatter plots and box plots are shown in each row as in Figure 2. **a)** shows we detected no significant difference in allele bias for *vit-2* reporter allele expression between control animals and *met-2(wam007)* animals; 0.000809 *met-2(wam007)* vs 0.000968 control, *P* = 0.266, Mann-Whitney Rank Sum Test, over 200 cells from each group, three independent experiments. **b)** shows we detected no significant difference in allele bias for *hsp-16.2* allele expression bias between control animals and *met-2(RNAi)* animals; 0.00311 met-2(RNAi) vs 0.0297 control, *P* = 0.595, Mann-Whitney Rank Sum Test, over 200 cells from each group, 3 independent experiments. **c)** shows we detected no difference in monoallelic expression of fluorescently tagged *idh-1* alleles between control animals and *met-2(RNAi)* animals; 0.00337 *met-2(RNAi)* vs of 0.00356 control, *P* = 0.730, Mann-Whitney Rank Sum Test, over 200 cells from each group, 3 independent experiments. **d)** shows we detected a significant difference in monoallelic expression of *eef-1A.1* reporter alleles between control and *met-2(RNAi)* animals. Variegated allele expression patterns can be seen in the images of animals undergoing *met-2(RNAi)* treatment, with some intestine cells showing virtually monoallelic mTagBFP2 or mNeptune expression. Control animals had a median intrinsic noise of 0.00254, whereas *met-2(RNAi)* animals had a significantly three fold greater median intrinsic noise of 0.00944; *P* < 0.001, Mann-Whitney Rank Sum Test, 180 total cells from each group from three independent experiments.

Next, we tested the heat shock inducible promoter from *hsp-16.2,* which controls expression of the small heat shock protein, HSP-16.2^49,50^. We conducted RNAi experiments as described above except we added a one-hour heat shock at 35°, 24 hours before we harvested animals for imaging. While *hsp-16.2* is sometimes expressed in a monoallelic fashion^32,35,36^, we found no significant difference in median intrinsic noise for *hsp-16.2* reporter alleles between control animals and animals on *met-2(RNAi)*, shown in Figure 3b.

Monoallelic expression of *IDH1* in tumors is associated with differences in survival time for patients with brain tumors^21^. We conducted RNAi experiments to examine the role of *met-2* in negatively regulating monoallelic expression for this gene, as above. We found no significant difference in median intrinsic noise/MAE for *idh-1* expression between *met-2* RNAi and EV control animals, shown in Figure 3c.

Finally, we tested strains that harbored reporter alleles controlled by the strong, constitutive promoter for *eef-1A.1,* a highly conserved translation elongation factor, also involved in the heat shock response^51^. When we placed these animals on *met-2(RNAi)* food, we saw a significant difference in median intrinsic noise compared to controls, shown in Figure 3d. In summary, *met-2* did not affect intrinsic noise for reporter alleles controlled by *vit-2* or *hsp-16.2* promoters, or endogenously tagged *idh-1*, but did affect intrinsic noise for alleles controlled by the *eef-1A.1* and *hsp-90* promoters. Thus, *met-2* negatively regulates MAE in a gene specific manner.

### MET-2 associated genes regulate MAE in unexpected ways

MET-2 is known to work with several other genes to regulate heterochromatin localization and silencing of genes, repetitive elements and transposons. Constitutive heterochromatin is tethered to the nuclear periphery by CEC-4 and LEM-2^52^. In worms, MET-2 and SET-25 are responsible for H3K9 methylation through two distinct regulatory pathways^40,41^. In a heterochromatin regulation pathway, MET-2 adds H3K9m1/2, and SET-25 adds H3K9me3^40^, with the help of LIN-61^41^. This pathway also requires LIN-65 to bind MET-2 in the cytoplasm and translocate it to the nucleus, and an additional MET-2 cofactor, ARLE-14, a conserved GTPase effector whose role is not completely understood^53^. In the gene and transposon silencing pathway, SET-25 directly adds H3K9me1/2/3 with the help of *nrde-3*^41^.

To determine the role of H3K9me HMTs and their associated cofactors in regulating MAE, we conducted RNAi knockdown of each gene in the *hsp-90* reporter allele strain and quantified intrinsic noise. Knockdown of *cec-4* and *lem-2* had no effect on MAE (Figure 4a&b). Knockdown of either *arle-14* or *lin-65* caused a significant increase in MAE compared to controls. The *arle-14(RNAi)* animals showed a subtler MAE phenotype (Figure 4c).The phenotype of *lin-65(RNAi)* animals was extreme (Figure 4d), and visually indistinguishable from *met-2(wam007)* and *met-2(RNAi)* animals (Figures 1&2). Thus, both *lin-65* and *arle-14* are negative regulators of MAE. Both *lin-61(RNAi)* and *set-25(RNAi)* showed the same BAE phenotype as controls (Figure 4e&f).

**Figure 4.**
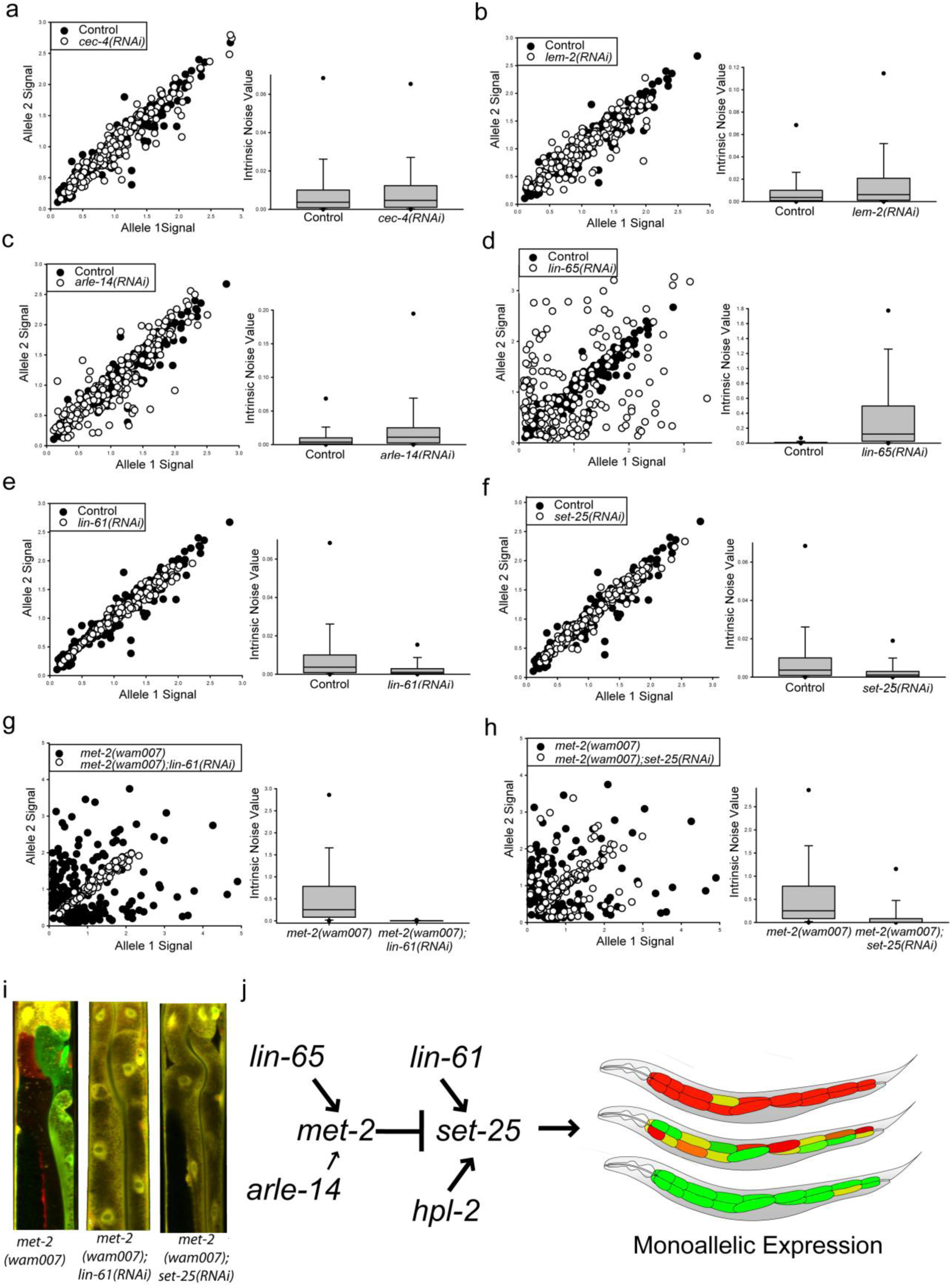
A unique genetic pathway regulates monoallelic expression. Scatter plots are shown in rows next to boxplots for **a-h**, as in Fig. 2 but without images. **a)** shows that cec-4 had no significant effect on MAE; *cec-4(RNAi)* 0.00467 vs control 0.00371 *P* > 0.05, Kruskal-Wallis One Way Analysis of Variance on Ranks followed by Dunn’s Method, >200 cells per group, three independent experiments. **b)** shows that lem-2 had no significant effect on MAE; *lem-2(RNAi)* 0.00624 vs control 0.00371 *P* > 0.05, Kruskal-Wallis One Way Analysis of Variance on Ranks followed by Dunn’s Method, >200 cells per group, three independent experiments. **c)** shows that *arle-14* negatively regulates MAE, with a significantly higher median intrinsic noise of 0.0110 compared to 0.00371 for control animals; *P* < 0.05, Kruskal-Wallis One Way Analysis of Variance on Ranks followed by Dunn’s Method, over 200 cells per group, three independent experiments. **d)** shows *lin-65* has a strong effect on negatively regulating MAE, with a significantly higher intrinsic of 0.0110 vs 0.00371, *P* < 0.05, Kruskal-Wallis One Way Analysis of Variance on Ranks followed by Dunn’s Method, over 200 cells per group, three independent experiments. **e)** shows that *lin-61* positively regulates MAE. *lin-61(RNAi)* animals had a significantly lower intrinsic noise of 0.00105 compared to control animals median intrinsic noise of median 0.00371; *P* < 0.05, Kruskal-Wallis One Way Analysis of Variance on Ranks followed by Dunn’s Method, over 200 cells per group, three independent experiments. **f)** shows that *set-25* positively regulates MAE. *set-25(RNAi)* animals had a significantly lower intrinsic noise of 0.00109 compared to control animals’ 0.00371; *P* < 0.05 for both comparisons, Kruskal-Wallis One Way Analysis of Variance on Ranks followed by Dunn’s Method, >200 cells per group, three independent experiments. **g)** shows that the enhanced MAE that happens in when MET-2 activity is eliminated is dependent upon LIN-61. The *met-2(wam007);lin-61(RNAi)* animals had a significantly lower intrinsic noise of 0.00160 than the *met-2(wam007)* animals’ median intrinsic noise of 0.254; *P* < 0.05 Kruskal-Wallis One Way Analysis of Variance on Ranks followed by Dunn’s Method for multiple comparison, >199 cells per group, three independent experiments. **h)** shows that the enhanced MAE that happens in when MET-2 activity is eliminated is dependent upon SET-25. The *met-2(wam007);set-25(RNAi)* animals had a significantly lower intrinsic noise of 0.00237 than the *met-2(wam007)* animals’ median intrinsic noise of 0.254; *P* < 0.05 Kruskal-Wallis One Way Analysis of Variance on Ranks followed by Dunn’s Method for multiple comparison, >199 cells per group, three independent experiments. **i)** shows images representative merged confocal microscope images of control, *met-2(wam007);lin-61(RNAi)*, and *met-2(wam007);set-25(RNAi)* animals’ “torso shots” – the anterior section of the intestine. **j)** shows the current genetic pathway controlling MAE in *C. elegans*. Additional statistical and genetic comparisons are shown in Supporting Information Section 2; see Supplemental Figure 3 for additional genetic tests.

To determine if loss of LIN-61 or SET-25 activity would suppress the increased MAE observed in *met-2(wam0007)* null animals, we quantified intrinsic noise for *hsp-90* reporter alleles in *met-2(wam0007)*;*lin-61(RNAi)* and *met-2(wam0007)*;*set-25(RNAi)* animals. Both double mutants had a significant decrease in MAE, demonstrating that the increase in MAE when MET-2 is genetically ablated is dependent on SET-25 and LIN-61, shown in Figures 4g&h. Figure 4i shows representative images of the double mutants and *met-2(wam007)* control animals.

In our initial screen for negative regulators of MAE, SET-25 had a coefficient of determination slightly higher than controls (Table 1). Therefore, we screened all other genes on Table 1 that had a coefficient of determination above controls (R^2^>0.928) for the ability to suppress the increased MAE in *met-2(wam007)* mutants. We identified one additional suppressor of MAE, *hpl-2*, a heterochromatin protein 1 homolog (Supplemental Figure 3 and 5). HPL-2/HP1 regulates chromatin, stress resistance and aging in *C. elegans*^54^. Taken together, these results show that *set-25* and *met-2* function antagonistically to regulate MAE, and the mechanism includes HPL-2 and LIN-61, which likely act as accessories to SET-25 function. Figure 4j shows the genetic pathway supported by our experiments.

### MET-2 and SET-25 require catalytic SET domains to regulate MAE

MET-2 and SET-25 are both H3K9 HMT proteins that use a catalytic SET domain to transfer methyl groups onto H3K9. In the context of germline immortality^55^, MET-2 also has an antagonistic relationship with SET-25 that does not require MET-2 HMT activity^43^. Because MET-2 has biological activity independent of its HMT SET domain, SET-25 and MET-2 may or may not require SET domains for regulating MAE.

To test the hypothesis that the regulation of MAE requires the SET domains of MET-2 and SET-25, we used CRISPR/Cas9 to generate single residue changes in the catalytic domains of *met-2* and *set-25,* and then measured MAE as described. We changed a critical cysteine residue to alanine, as was previously demonstrated to effectively nullify SET domain activity for *met-2*^43^ (Figure 5a-c). The resulting mutations were *met-2(wam406)*, which reprograms MET-2 for a C1237A residue change, and *set-25(wam404)*, which reprograms SET-25 for a C645A residue change. Additionally, we made a third strain with both *met-2(wam406)* and *set-25(wam404)* mutations.

**Figure 5.**
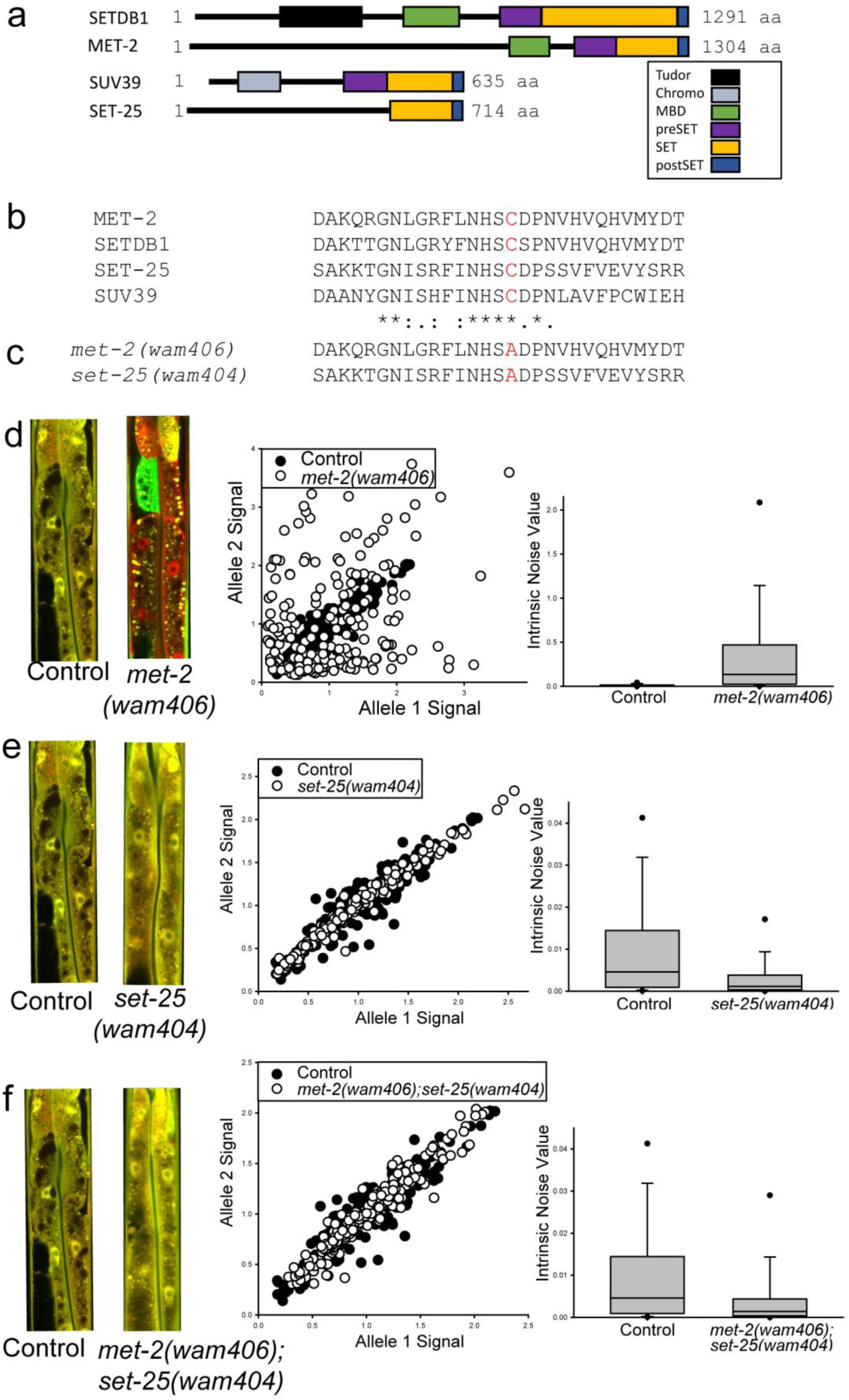
Cysteine to alanine mutation in the SET domain of SET-25 or MET-2 causes loss of MAE-regulating activity. Rows **e-g** show images, scatter plots and boxplots as described in Figure 2. **a)** shows graphical alignment of human SETDB1 and SUV39 with worm homologs. Predicted domains are highlighted by different colored boxes. **b)** shows a 2D image of the predicted 3D structure of SET-25 (left panel) and a zoomed in view of the critical cysteine residue in the SET domain highlighted red (right panel), which we reprogrammed to alanine. **c)** shows the text alignment of residues in the SET domains of human proteins and their worm homologs, highlighting the critical cysteine residue in red. **d)** shows the text alignment of the SET domain of *met-2(wam406)* and *set-25(wam404)* mutants we edited the genomes of to reprogram expression of cysteine in **c** to alanine, highlighted in red. **e)** shows that the *met-2* SET domain mutant we created here phenocopies the null mutant, having significantly higher intrinsic noise than control animals; 0.134 for *met-2(wam406)* vs. 0.00459 in controls, *P* < 0.05, Kruskal-Wallis One Way Analysis of Variance on Ranks followed by Dunn’s Method for Multiple Comparison, >200 cells per group, three independent experiments. **f)** shows that the set-25 SET domain mutant we made phenocopies the set-25(RNAi), with significantly lower intrinsic noise than controls; 0.00114 in *set-25(wam404)* vs 0.00459 for control, *P* < 0.05, Kruskal-Wallis One Way Analysis of Variance on Ranks followed by Dunn’s Method for Multiple Comparison, >200 cells per group, three independent experiments. **g)** shows that *met-2;set-25* double SET-mutant animals have significantly lower intrinsic noise than *met-2(wam406)* animals and control animals; 0.00140 for *met-2(wam406);set-25(wam404)* vs. 0.00459 for control or 0.134 for met-2(wam406), *P* < 0.05 for both comparisons, Kruskal-Wallis One Way Analysis of Variance on Ranks followed by Dunn’s Method for Multiple Comparison, >200 cells per group, three independent experiments.

We found the *met-2(wam406)* animals phenocopied the *met-2(RNAi)* and *met-2(wam007)* null animals, with all animals showing extreme MAE and higher intrinsic noise compared to controls (Figure 5d). Next, we measured allele expression in *set-25(wam404)* animals and found this mutation phenocopied the *set-25(RNAi)*, showing significantly less MAE (lower intrinsic noise/more BAE) compared to controls, shown in Figure 5e. Finally, we measured the intrinsic noise in the *met-2(wam406);set-25(wam404)* double point mutation mutants, and found they had significantly lower intrinsic noise/MAE than both control animals and *met-2(wam406)* animals (Figure 5f); these animals showed a nearly complete biallelic expression pattern. These results show that the antagonistic regulation of MAE by MET-2 and SET-25 requires the H3K9 HMT catalytic SET domains of both proteins to be intact.

### MET-2 and SET-25 antagonistically regulate germline immortality and aging

Like monoallelic expression (Figure 4g&h), germline immortality has been shown to be antagonistically regulated by *set-25* and *met-2*^43^. In those studies, the *met-2* catalytic domain mutants phenocopied the *met-2* null^43^, causing a mortal germline phenotype. However, the role of the SET domain of SET-25 in germline immortality was unknown, and those experiments were conducted at a higher temperature of 25°^43^. In order to determine if the SET domain of SET-25 is required to kill the germline in *met-2* mutants, we conducted fecundity experiments at 20°, where *set-25*;*met-2* mutants have fewer developmental difficulties compared to 25°^56^. Here, we analyzed hermaphrodite self-progeny production over the first four days of adulthood at 20°^57^ for *met-2(wam007)*, *met-2(wam406), set-25(wam404), and met-2(wam406);set-25(wam404)* animals.

We found the control animals and *set-25(wam404)* animals to be generally similar in progeny production (Figure 6a-f). However, one measure distinguished control and *set-25(wam404)* animals. The *set-25(wam404)* mutants actually had higher than control progeny production on the fourth day of adulthood (Figure 6d), though we did not detect a difference in total progeny production (Figure 6e) or interindividual variation in progeny production (Figure 6f).

**Figure 6.**
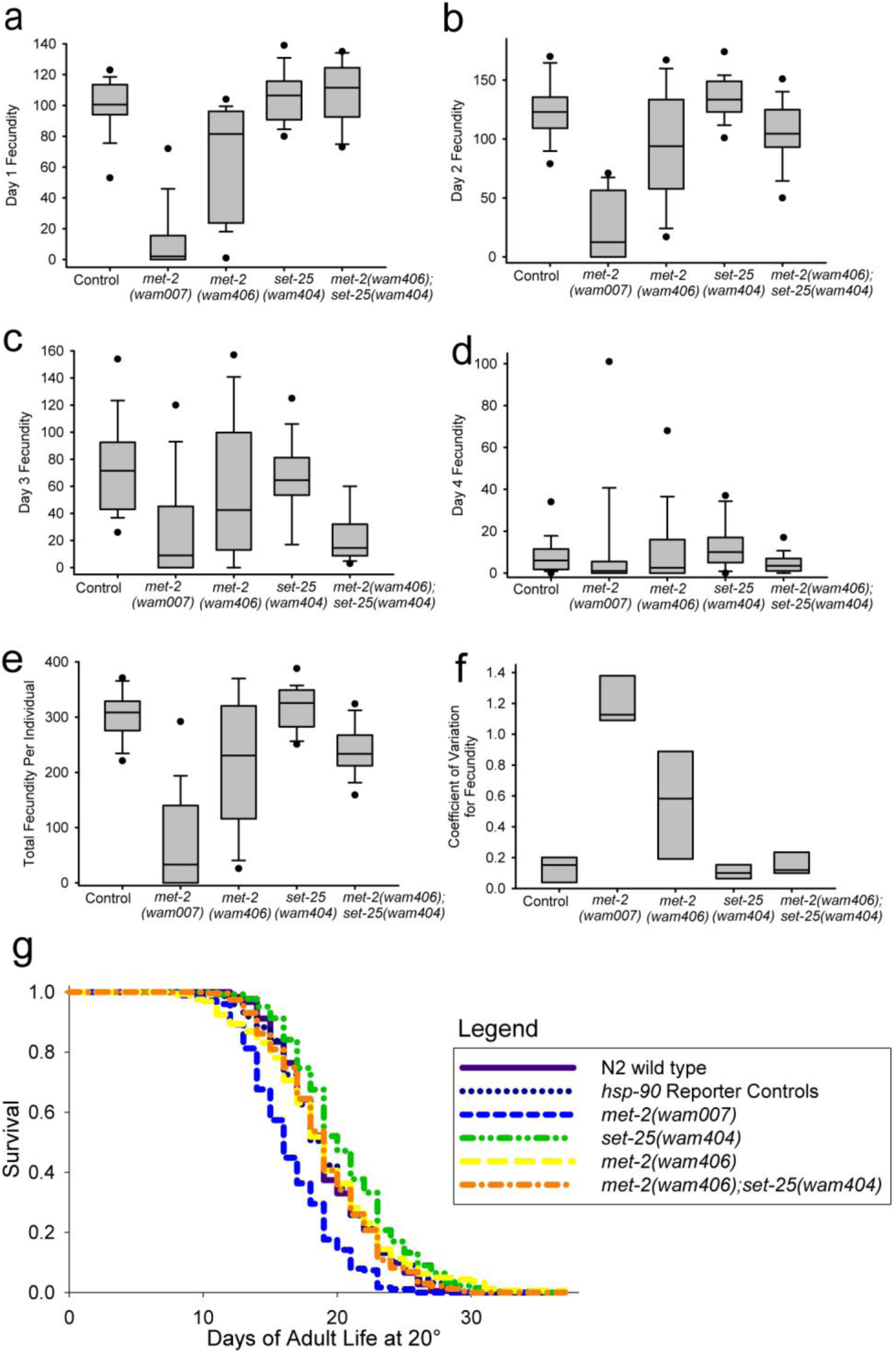
The antagonistic relationship between MET-2 and SET-25 also regulates germline mortality and aging. Boxplots of daily fecundity are shown in **a-d**. Boxplots show total fecundity for days 1-4 of adulthood in **e**, with interindividual variation in fecundity plotted in **f**. **a)** shows progeny production on day 1 of adulthood. The *met-2* mutants had significantly lower progeny production than control animals. The *met-2(wam406);set-25(wam404)* animals had significantly higher progeny production than the *met-2(wam406)* animals on this day; *P* < 0.05 for noted comparisons, Tukey Test, three independent experiments with six individuals per group per experiment. **b)** shows progeny production on day 2 of adulthood. The *met-2* mutants still had significantly lower progeny production than wild type; *P* < 0.05 for both comparisons, Tukey Test, three independent experiments with six individuals per group per experiment. **c)** shows day 3 fecundity, with the same trend for *met-2* mutants persisting. The *met-2(wam406);set-25(wam404)* animals had significantly higher progeny production than the *met-2(wam406)* animals on this day; *P* < 0.05 for noted comparisons, Tukey Test, three independent experiments with six individuals per group per experiment. **d)** shows day 4 fecundity with almost no difference between groups. Interestingly, the *set-25* mutants had significantly more progeny than all other groups on day 4 of adulthood; **e)** shows total fecundity for each group. Total fecundity was significantly lower than control animals for both *met-2* mutants; *P* < 0.05 for both comparisons, Student-Newman-Keuls Method, three independent experiments with six individuals per group per experiment. For total progeny, the double catalytic mutant, *met-2(wam406)*;s*et-25(wam404)* had significantly more progeny than the *met-2* knockout, *met-2(wam007)*, but not significantly more progeny than the *met-2* SET domain mutant, *met-2(wam406),* and significantly less than control and *set-25(wam404)* animals; *P* < 0.05, Student-Newman-Keuls Method, three experiments per group, six individuals per group per experiment. **f)** shows boxplots of the coefficient of variations from the individual experiments. There was significantly more interindividual variation in fecundity for the *met-2* mutants compared to control animals, as measured by the Coefficient of Variation (CV); *P* < 0.05 for both comparisons of CV, Holm-Sidak method, three experiments with three CV values per group. **g)** Lifespan line plots of different groups of animals are shown. X axis shows the days of adult life and the y axis shows the fraction of the population surviving. We did not detect a difference in median lifespan between N2 control animals median lifespan of 19 days and any other group besides *met-2(wam007)* animals, with a median lifespan of 16 days, and *set-25(wam404)* animals, with a median lifespan of 20 days; *P* < 0.05 for all comparrisons, Kruskal-Wallis One Way Analysis of Variance on Ranks followed by Dunns Method, N of animals per group ranged from 160-265. Additional statistical and graphical comparisons are shown in Supporting Information Section 2; see Supplemental Figure 4.

Generally, both *met-2(wam007)* and *met-2(wam406)* showed a mortal germline phenotype at 20° (Figure 6a-e). The *met-2(wam406)* worms were not as severely sterile as the *met-2(wam007)* (Fig. 6e), as has been seen before at 25°^43^. The *met-2(wam406);set-25(wam404)* double SET domain mutants did not fully rescue the reduced fertility caused by *met-2(wam406)*, nor did we detect a change in total fertility between *met-2(wam406)* and the *met-2(wam406);set-25(wam404)* double mutants (Fig. 6a-e; Supplemental Figure 4). However the double mutants were distinct from both *met-2* mutants in two ways. First, the double mutants show a delayed defect in progeny production. These animals produce a normal number of progeny on day 1 of adulthood (Fig. 6a), with a small decrease on day 2 (Fig. 6b), and a precipitous decrease in progeny on day 3 of adulthood (Figure 6c). This is different from the mortal germline phenotypes of both *met-2* mutants, where progeny is consistently down for *most* individuals on the first three days of production (Figures 6a-c). Second, the interindividual variation in progeny production of both *met-2* mutants was significantly higher than control animals (Figure 6f). This difference in interindividual variation distinguishes the phenotypes of the *met-2(wam406)* and the *met-2(wam406);set-25(wam404)* animals, which both had lower than control progeny production. This increased interindividual variation in progeny production of *met-2* mutants (Figure 6f) is suggestive of monoallelic expression being a form of epigenetic gambling^58^, as some individuals in the *met-2* mutant populations had abundant progeny; they were not all low fecundity.

We quantified lifespan for the same animals in which we quantified progeny production. We found that, compared to control animals, the *met-2(wam007)* animals had a significantly shorter lifespan, whereas the *met-2(wam404)* animals, which harbor the C to A residue change in the catalytic SET domain, had normal lifespans (Figure 6g). Conversely, the set*-25(wam404)* animals, which harbor the same C to A residue change, had an increased lifespan (Figure 6g). We conclude that *met-2* and *set-25* antagonistically control lifespan. Importantly, and unlike its effects on germline mortality and MAE, the shortened lifespan seen in *met-2* null animals is distinct from the *met-2(wam406)* mutation, which had wild type lifespan.

### Somatic RNAi initiation and inheritance is unaffected in animals lacking MET-2 and SET-25 HMT activity

The MAE regulators we identified could be involved with RNAi or inheritance of RNAi in the soma because both processes involve silencing at the chromatin level^59^. *C. elegans* can use RNAi to both alter and inherit states of gene expression in the germline^60^. Previous research has shown that loss of *met-2* causes a permanent, transgenerational inheritance of RNAi-dependent silencing for at least two genes expressed in the germline^60,61^. Therefore, *met-2* could affect somatic RNAi, or inheritance of it. Additionally, *set-25* could affect initiation or inheritance of RNAi, because it affects heritable differences in chaperone expression levels^62,63^. Thus, we wanted to determine if the somatic RNAi silencing machinery was functional, and if somatic RNAi inheritance would be altered, in our *met-2* and *set-25* mutants.

To determine if initiation or inheritance of RNAi was affected by MAE regulators, we tested their ability to perform somatic RNAi and inherit those RNAi states. We tested the effects of *met-2(wam007)*, *met-2(wam406), set-25(wam404), and met-2(wam406);set-25(wam404)* on the ability of animals to both initiate and inherit RNAi (Figure 7). We used mCherry protein signal as a measure of the effectiveness of the RNAi treatment. After one generation on *mCherry(RNAi)*, all of the strains showed typical RNAi kinetics, with the near complete loss of mCherry in all tissues except the pharynx, which we and others^64^ find to be resistant to RNAi. After a second generation on RNAi, all mCherry signal (except for some expression in the pharynx) was gone from all strains. When we switched the animals back to OP50 for a single generation, mCherry signal returned in all strains. After two generations on OP50, all of the strains showed mCherry signal similar to controls on EV. These results show that the loss of MET-2 protein or HMT catalytic activity, or loss of or SET-25 HMT catalytic activity have no detectable effects on the initiation or inheritance of somatic RNAi.

**Figure 7.**
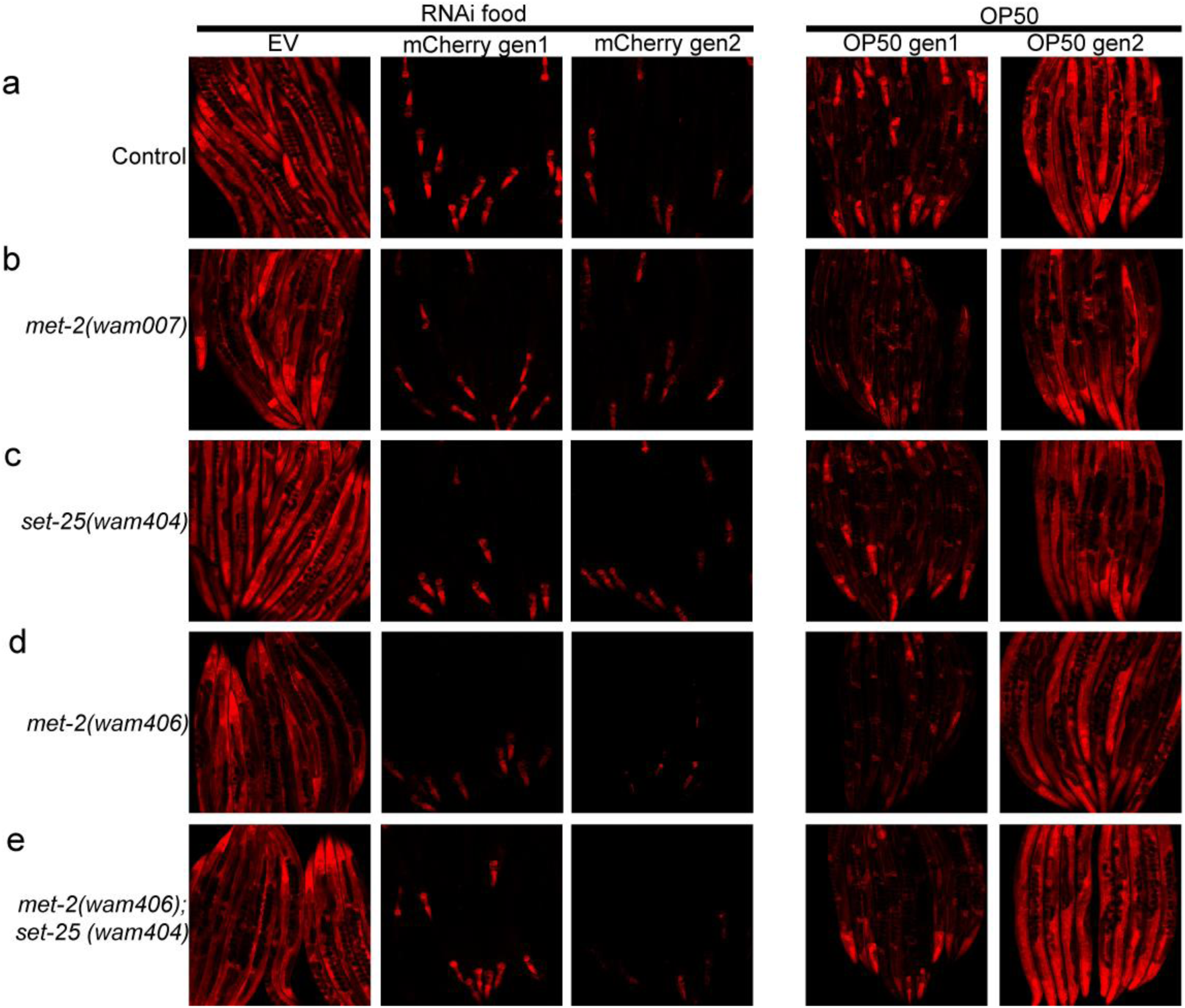
Somatic RNAi initiation and inheritance are not affected by MAE regulators. Each row (**a-e**) shows a different genotype. Each column is a different generation. Images show the loss of mCherry signal over two generations of RNAi, and the reemergence of mCherry signal after two generations off of RNAi food. None of the conditions we tested had any differences in the initiation or stopping of RNAi. Images are representative of results from three independent experiments.

### MAE is not heritable

In cell culture, MAE persists across mitotic divisions, which presumably means that it persists throughout the lifetime of the cell^9,20,31^. In intact *C. elegans*, we observe that MAE is persistent in adults, and sometimes mitotically propagated throughout a tissue (e.g., the intestine), especially in *met-2* mutants. Additionally, *met-2* and *set-25* were associated with inheritance of silencing/gene expression levels in the germline^60,61^ and soma^63^, respectively.

To determine if the increased MAE we describe here is heritable, we imaged day 2 adult *met-2(wam406)* animals (second day of adulthood for *C. elegans* hermaphrodites at 20°) that would act as the P0 parents, put the animals back in culture to lay eggs, and then imaged the F1 progeny from each individual parent. An example of these experiments is show in Figure 8. The patterns of MAE between parent and offspring, and between siblings, are different. For example, a mostly variegated parent can have offspring that have fully monoallelic tissues, such as the intestine, and the expressed allele can be either paternal (green) or maternal (red). We conclude from these experiments that MAE is not heritable.

**Figure 8.**
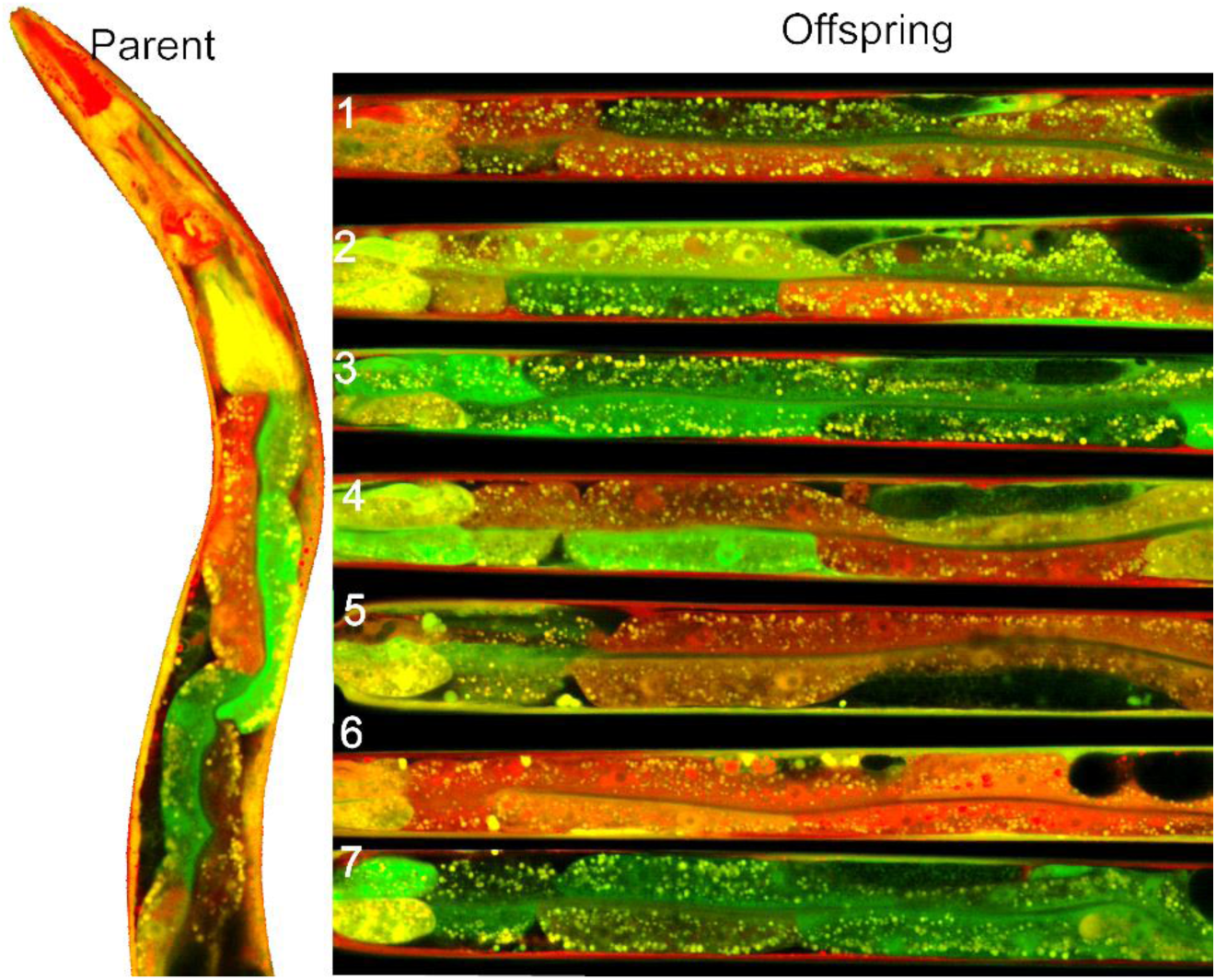
MAE is not heritable. Image on left is a *met-2(wam406)* mutant parent hermaphrodite expressing differently colored *hsp-90* promoter controlled fluorescent alleles. Images on right are seven individual heterozygous progeny of the heterozygous parent. The heterozygous parent has a variegated pattern of expression and produces an array of diverse progeny with both variegated patterns and virtually monoallelic patterns of allele expression in the intestines. Results are representative of three independent experiments.

### Maternal MET-2 and SET-25 act in the early embryo to control MAE

In *C. elegans*, the entire set of 20 intestine cells are descended from the single E-cell in the 8-cell embryo. Because we see persistent MAE throughout the entire intestine in some animals, it suggests that MAE can be initiated in the E-cell, propagated throughout mitotic divisions, and maintained into adulthood (e.g., Figs. 1a,d). In the 8-cell embryo, there is very tight control of embryonic gene transcription^65^. We hypothesized that because of the early initiation of MAE, we (here), and others^66^ observe, and the rapid pace of *C. elegans* development, maternal or paternal proteins are responsible for initiation of MAE in the E-cell.

To test our hypothesis, we performed two sets of reciprocal crosses. First, we crossed *met-2wam(406)*;*P_hsp-90_::mEGFP* males with *P_hsp-90_::*mCherry hermaphrodites, and performed the reciprocal cross of *P_hsp-90_::mEGFP* males with *met-2(wam406)*; *P_hsp-90_::* mCherry hermaphrodites (Figure 9a&b). When *met-2(wam406)* was crossed in from the male, all progeny showed BAE (Figure 9c), which is expected in a mutant that is heterozygous recessive. However, when *met-2(wam406)* was crossed in from the female, all progeny showed MAE (Figure 9d), despite being heterozygous for a recessive *met-2* mutation. The increase in median intrinsic noise when animals receive *met-2(wam406)* from the female germline was highly significant, shown in Figures 9e&f.

**Figure 9.**
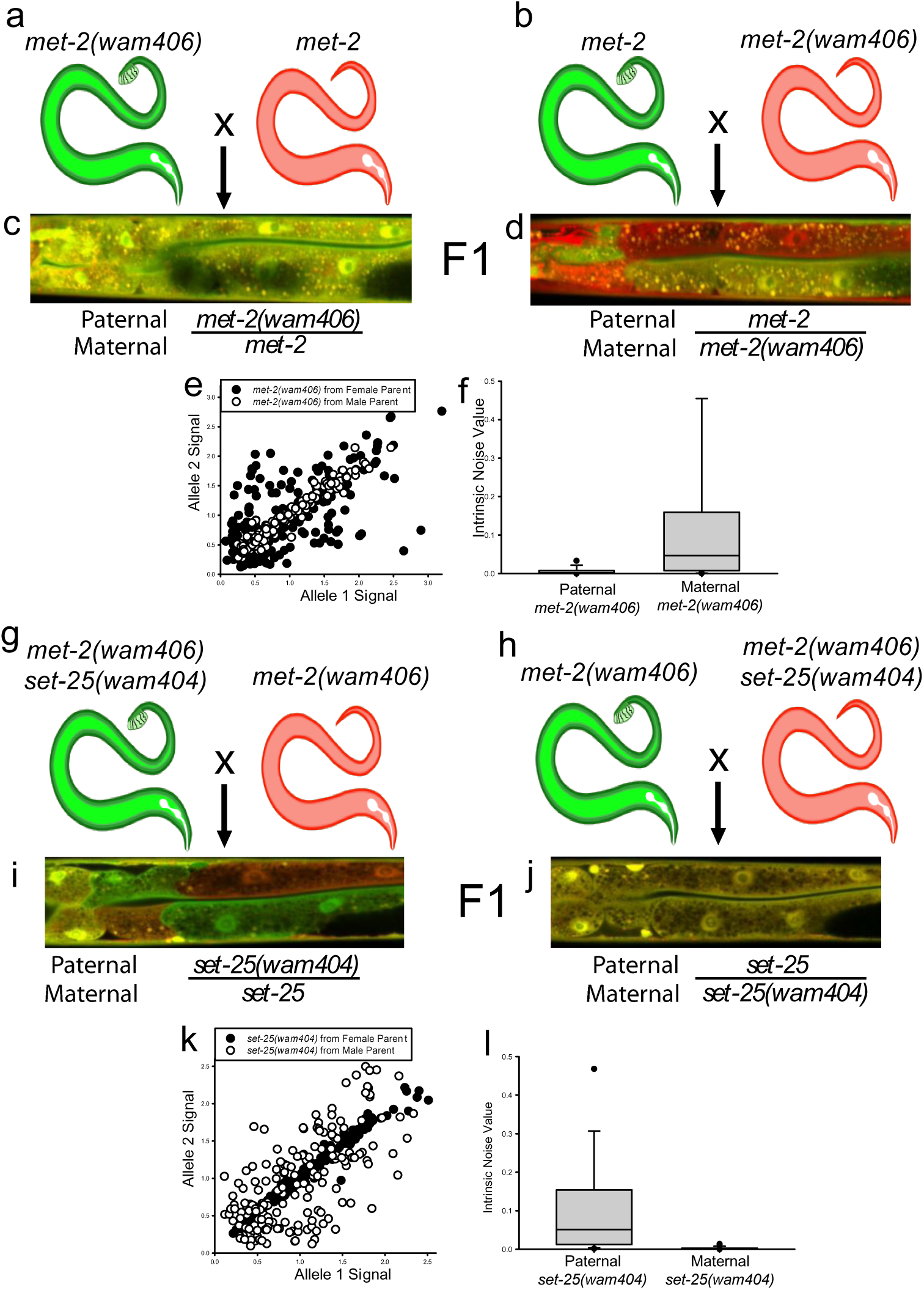
Maternal MET-2 and SET-25 act in the early embryo to regulate MAE. **a)** and **b)** show reciprocal crosses generating F1 hybrids heterozygous for recessive *met-2(wam406),* which results in MAE when homozygous. F1 hybrids are shown imaged for *hsp-90* reporter allele expression in **c)** and **d)** respectively. **c)** shows a typical F1 hermaphrodite heterozygous for *met-2(wam406)*, bred in from the paternal sperm. **d)** shows a typical F1 hermaphrodite heterozygous for *met-2(wam406)*, bred in from the maternal oocyte. **e)** shows scatterplots of cells from F1 animals from each reciprocal cross shown in **a&b**. **f)** shows boxplots of intrinsic noise of cells from F1 animals fromeach reciprocal cross shown in **a&b**. The F1 *met-2(wam406)* heterozygotes that got the *met-2(wam406)* allele from their mother had the same increased intrinsic noise as homozygous *met-2* mutants, with a significantly higher intrinsic noise than animals that got the mutant *met-2* from their father; 0.0469 median intrinsic noise for maternal *met-2* vs 0.00193 for paternal *met-2*, *P* < 0.001, Mann-Whitney Rank Sum Test, N=180 cells per group, three independent experiments. **g)** and **h)** show reciprocal crosses generating F1 hybrids homozygous for the *met-2(wam406)* mutation, which causes MAE, and heterozygous for recessive *set-25(wam404),* which results in BAE when homozygous. F1 hybrids are shown imaged for *hsp-90* reporter allele expression in **i)** and **j)** respectively. **i)** shows a typical F1 hermaphrodite heterozygous for *set-25(wam404)*, bred in from the paternal sperm. **j)** shows a typical F1 hermaphrodite heterozygous for s*et-25(wam404)*, bred in from the maternal oocyte. **k)** shows scatterplots of cells from F1 animals from each reciprocal cross shown in **g&h**. **l)** shows boxplots of intrinsic noise of cells from F1 animals from each reciprocal cross shown in **g&h**. The F1 *met-2(wam406)* homozygous, *set-25(wam404)* heterozygous animals that got the *set-25(wam404)* allele from their mother had the same decreased intrinsic noise as homozygous *set-25* mutants, with a significantly lower intrinsic noise than animals that got the mutant *set-25(wam404)* allele from their father; 0.00109 median intrinsic noise for maternal mutant *set-25* vs. 0.0508 for paternal mutant *set-25*, *P* < 0.001, Mann-Whitney Rank Sum Test, N= 180 cells per group, three independent experiments.

Next, we performed a cross of *met-2(wam406);set-25(wam404);P_hsp-90_::mEGFP* males with *met-2(wam406);P_hsp-90_::mCherry* hermaphrodites (Figure 9g), and the reciprocal cross of *met-2(wam406)*;*P_hsp-90_::mEGFP* males with *met-2(wam406);set-25(wam404);P_hsp-90_::mCherry* hermaphrodites (Figure 9h). The F1 animals homozygous for *met-2(wam406)*, and heterozygous for *set-25(wam404)* received from the father, showed the expected MAE (Figure 9i). This is the Mendelian phenotype that would be expected for a recessive heterozygous mutation in a positive regulator of MAE, in a homozygous negative MAE regulator background. The F1 progeny homozygous for *met-2(wam406)* and heterozygous for *set-25(wam404)* from the mother showed unexpected BAE (Figure 9j). This result is inconsistent with the expected Mendelian phenotype for a heterozygous recessive mutation in a positive regulator of MAE, in a negative MAE regulator background. The median intrinsic noise for F1 progeny heterozygous for *set-25(wam404)* was significantly greater for animals that received the mutant *set-25* from their mother, compared to animals that received the mutant *set-25* from their father, shown in Figure 9k&l. These results show that MAE is regulated in the early embryo by the catalytic SET domain activity of maternal MET-2 and SET-25. A model based on our findings is shown in Figure 10.

**Figure 10.**
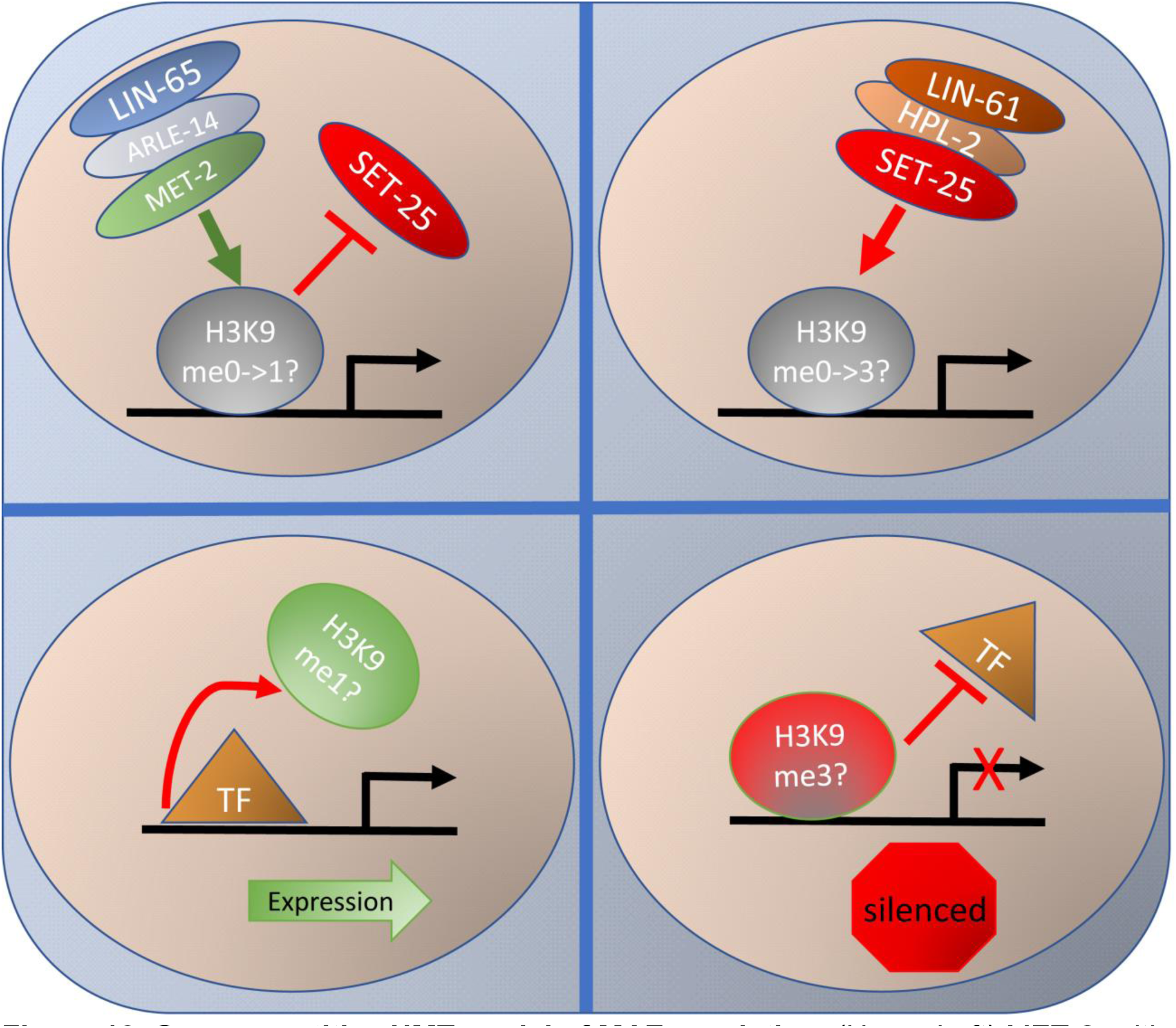
Our competitive HMT model of MAE regulation. (Upper Left) MET-2, with the help of ARLE-14 and LIN-65, binds an allele near the promoter, allowing MET-2 to methylate H3K9, and block SET-25 binding. (Lower Left) This results in transcription factor (TF) binding and expression of that allele. (Upper Right) If SET-25 interacts with an allele before MET-2, it can trimethylate H3K9. (Lower Right) This results in blocking transcription factor binding and the silencing of an allele. Promoters have different affinities for one or both HMTs to cause different patterns of expression in a probabilistic fashion.

## DISCUSSION

### Our model for regulation of monoallelic expression

We report a working model for a developmental genetic pathway that controls MAE in the worm soma. It consists of two known H3K9 HMTs working antagonistically along with known associated proteins. Our data suggests that HPL-2 and LIN-61^41^ enable maternal SET-25 to bind to histones associated with promoters of genes with MAE potential, and for intestine tissue, this happens in the E-cell of the 8-cell embryo (Figs. 4,9&10). In parallel, the MAE-related silencing activities of HPL-2, LIN-61 and SET-25 are inhibited by LIN-65, MET-2, and to a lesser extent ARLE-14. LIN-65 translocates MET-2 to the nucleus^53^, where it is helped by ARLE-14^53^ to act as a transcriptional-silencing-repressor of SET-25, affecting the timing of initiation of SET-25 based silencing to generate different patterns of silencing (Figs. 4,9&10 and Supplemental Figure 1).

In the context of heterochromatin^40^ and transposons^41^, MET-2 causes silencing. Here, we show that MET-2 is acting to inhibit silencing. Our experiments strongly suggest MET-2 inhibits SET-25 using histone methylation, because when MET-2 HMT activity is disrupted (Figs 5&9), SET-25 uses its own HMT activity to silence alleles, leading to MAE (Figure 5). When SET-25 lacks HMT activity, it cannot silence alleles.

If MET-2 uses its catalytic SET domain to inhibit SET-25 activity, it is most likely depositing H3K9 markings, based on our current understanding of MET-2 activities. SET-25 can methylate H3K9 independently of MET-2^40–42^. If SET-25 HMT activity is inhibited by MET-2 HMT activity, then there are two plausible scenarios for probabilistic regulation of non-heritable MAE. One scenario is that SET-25 cannot initiate methylation on H3K9me1. In this scenario, H3K9me1 put down by MET-2 leads to expression of an allele and inhibition of SET-25 HMT activity. However, if MET-2 marks histones with H3K9me2, or if MET-2 is inhibited from methylating H3K9 altogether, then SET-25 can add the third methyl to H3K9me2, or it could add two or three methyl groups to H3K9me0 to inhibit expression of an allele. In this scenario, SET-25 cannot act on H3K9me1 or it can do so with low efficiency. This model could explain an antagonistic relationship wherein both MET-2 and SET-25 require their SET domains for H3K9 methylation at MAE loci.

In another scenario, SET-25 and MET-2 could be working with other yet unidentified proteins to regulate MAE via H3K9 methylation states. Alternatively, SET-25 and MET-2 could have additional SET-related activities that modify other residues, perhaps being part of a combinatorial code^14,15^ or even new marks^67^. ChIP based studies in mammalian tissue found the H3K9me3 marking to be non-informative for ChIP based identification of MAE^14,15^, consistent with a non H3K9me role for MET-2 and SET-25. Yet, in other biological contexts, H3K9 marks are the critical determinants of monoallelic expression^20^.

Regulation of MAE is gene specific, with, for example, DNA methylation controlling MAE for some genes^12^ and H3K9 markings controlling MAE for others^20^. So, it is not entirely surprising that *hsp-90* and *hsp-16.2* reporter alleles are regulated differently. That is, both *hsp-90* and *hsp-16.2* have monoallelic potential, but only *hsp-90* responded to perturbation of *met-2*. In worms, like in humans^12,20^, there are multiple regulatory pathways controlling monoallelic expression, and we have yet to define the *hsp-16.2* MAE-regulating pathway. Additionally, there may be other yet unidentified accessory proteins that attract MET-2 and SET-25 proteins to specific MAE promoters; they could be essential proteins that are not detectable with genetics. Future studies will elucidate *met-2* independent pathways regulating MAE. Moreover, technologies such as Fiber-seq^68^, that can be used to examine nucleosome occupancy changes at single molecule resolution, in conjunction with genetic studies, will help define additional MAE control pathways.

### The genetic control pathway for MAE is unique, but overlaps with heterochromatin, transposon silencing and germline immortality pathways

In addition to regulating monoallelic expression, *met-2/SETDB1* and *set-25/SUV39/G9a* also regulate germline immortality^43,55^, transposon silencing^41,69^, and heterochromatin^40,52,53,70^. Here we found that the SET domain of SET-25/SUV39 was required for negative regulation of germline immortality in *met-2(wam406)* C695A mutants. While genes like *hrde-1* are reported to also negatively regulate germline immortality in *met-2* mutants^61^, the knockdown of *hrde-1* had no effect on monoallelic expression (Table 1 and Supplemental Figure 3).

Germline immortality and somatic monoallelic expression could be physiologically connected. When *met-2* HMT activity is ablated via the SET domain mutation, the output of the germline diminishes on average. Interestingly, the variation is progeny number between *met-2* HMT activity mutants is more variable than controls. This is consistent with the idea that random silencing of alleles by SET-25 might act to randomly alter investments between the soma and germline^71^. Under laboratory conditions, SET-25 is acting to slightly restrict lifespan and fertility, and MET-2 is acting to promote lifespan and fertility (Figure 6). If MAE and progeny production are linked, then it is possible that MAE, as a form of epigenetic gambling^58^, could be selected for, via selection for or against pro or anti germline factors in different biological scenarios. A population without this dynamic system that antagonistically controls aging (often linked to stress resistance^72^), fertility and somatic monoallelic expression might not be as fit to survive environmental uncertainties. This sort of intangible epigenomic product created by the MET-2/SET-25 MAE control system has been speculated to be active in the early embryo^7^, consistent with what we observed here (Figure 9) and what the armadillo monozygotic quadruplet sequencing data indicates^66^.

Prior work has shown that SET-25 regulates heterochromatin and silences the expression of genes and transposons via two different transposon control pathways^41^. The first is the MET-2-LIN-61-SET-25 heterochromatin control pathway in which SET-25 works cooperatively with LIN-61 and MET-2 to trimethylate histones in heterochromatin and at transposon loci^41^. In the second control pathway, SET-25 functions with NRDE-3 to add all three methyl groups to H3K9. Here we found SET-25 and LIN-61 promote the silencing of alleles in the absence of MET-2 (Fig. 4). The function of LIN-61 in regulating MAE is not completely clear, but it may be generally assisting SET-25 to bind H3K9^73^. Similarly, we detected no effect of *nrde-3(RNAi)* on MAE in our initial screen or in subsequent experiments (see Table 1 and Supplemental Figure 2). Thus, the current six-gene network performing MAE regulation in *C. elegans* overlaps with known genetic pathways regulating other silencing-related phenomenon, but seems unique in its utilization of genetic components.

Here, we have worked toward an understanding of the play-by-play events regulating the initiation, propagation and maintenance of monoallelic expression^8^. We discovered that two conserved H3K9 HMTs and their accessory proteins antagonistically control MAE in the early embryo. This developmental genetic pathway can explain how monoallelic expression is initiated and propagated throughout a tissue. The homologs of genes we identified here may regulate MAE in mammalian systems, where H3K9 methylation is known to control MAE-related silencing^20^. This model system can be used to identify additional MAE regulatory pathways and corresponding changes to chromatin. Further investigations will enable us to understand the gene-specific mechanisms that control random autosomal monoallelic expression, which can be developed into technologies to affect positive change in human health.

## METHODS

### Strain Creation

We used alt-R CRISPR/Cas9 (IDT, Newark, NJ) to generate endogenous gene tags and point mutations.^13^ For making single point mutations in the *met-2* and *set-25* catalytic domains, and for making the met-2 knockout, we used ssODNs as repair templates (IDT, Coralville, IA). For endogenously tagging *idh-1*, we used PCR products as repair templates. CRISPR target and repair template sequences are listed in Supplementary Table 3. Primers are listed in Supplementary Table 2. CRISPR point mutations were made directly into the heterozygotic GFP/mCherry reporter allele expressing strains. Genomic DNA was harvested from each CRISPR edit and a region spanning the repair template was amplified by PCR and sequenced (Genewiz, Burlington, MA). The resulting strain names and genomic insertion designations are shown in Supplementary Table 1.

### Animal Husbandry and RNAi

Animals were cultured as previously described^74^. Briefly, we maintained all strains in 10cm petri dishes on NGM seeded with OP50 *E. coli* in an incubator at 20°. All experiments were performed at 20°, with the exceptions of a heat shock experiment series and male generating procedures noted below. RNAi experiments used NGM supplemented with 50ug/ml carbenicillin, 10 ug/ul tetracycline, and 1mM IPTG. All strains used in this study are listed in Supplementary Table 1. A table of crosses can be found in Supplementary Table 4. To generate heterozygous GFP/mCherry expressing strains, we generated GFP expressing males by subjecting 20 L4s to a 30° heat shock for 5-6 hours. For all strains, the GFP allele was introduced through the male germline. We screened for heterozygous animals that express both GFP and mCherry on a fluorescence stereoscope. We maintained heterozygotes by picking them away from homozygous animals and onto fresh, OP50-seeded NGM growth plates each generation. We performed experiments on heterozygous animals that were at least five generations beyond the initial cross to avoid paternal allele expression bias. To synchronize animals for experiments, we conducted two hour egg lays onto 10cm NGM or RNAi plates (10 heterozygous animals per plate). For experiments with heat shock to quantify *hsp-16.2* reporter expression, we performed a one-hour heat shock at 35° on one day old adult animals by placing animals on their NGM growth plates into a 35° incubator for one hour and then returning them back to the 20° incubator until imaging the next day (24 hours post heat shock).

For RNAi experiments, day 2 adult worms heterozygous for reporter alleles were placed on RNAi food or EV to lay eggs for two hours before the parents were removed. The process was repeated on day 4 for a total of two generations on RNAi food. On day eight from the start of the experiment, synchronous day 2 adults were harvested for microscopy.

### Progeny Production Per Individual

To quantify differences in the fertility of the self-fertile *C. elegans* hermaphrodites, we picked L4 animals onto single plates and allowed individual animals to lay eggs for the first four days of adulthood at 20°. P0 worms were moved to new plates each day. We quantified progeny per individual per day by counting the number of hatched progeny at the L4/YA age, two days after moving the hermaphrodite. Progeny for all worms were measured approximately 20 generations after the worms were created.

### Lifespan Measurements

To quantify genetic effects on aging, we quantified lifespan using the semi-automatic WormBot system^75^. Briefly, L4 animals are placed in groups of approximately 20-30 onto OP50 seeded NGM media supplemented with 50uM FUDR to inhibit progeny production in 12-well plates with clear plastic bottoms and lids. The WormBot system captures images/movies of each well that we then manually review for the cessation of movement/death.

### RNAi kinetics in mutant strains

To determine if the somatic RNAi machinery was functional and if RNAi heritability was altered in *met-2* and *set-25* mutants, strains homozygous for *P_hsp-90_::mCherry* reporters were grown on OP-50 food for at least five generations before conducting RNAi. For egg lays, 10 gravid adults were placed on RNAi food or OP50 for two hours to lay eggs before removing the P0s. On experiment days 4 and 8, egg lays were conducted on RNAi food. On experiment days 12 and 16, egg lays were conducted on OP50 food.

After each egg lay, the remaining worms from each plate were harvested for imaging. To image, 15-20 worms from each plate were anesthetized on agarose slides and imaged on a Zeiss LSM780 confocal microscope with a 10x air objective. All RNAi clones were sequenced prior to seeding NGM plates.

### Heritability Experiments

To determine if patterns of monoallelic expression were heritable, we used a Leica M165F Fluorescent stereoscope with a 500nm long-pass emission filter to identify day 2 adult *met-2(wam406)* individuals that were heterozygous for fluorescent red and green *hsp-90* reporter alleles. We picked individuals with either variegated or monoallelic expression patterns in their intestine onto individual plates. We let each individual animal lay eggs on a 6cm NGM plate for four hours, then we mounted each individual animal on an agarose pad to record an image by confocal microscopy as we describe below.

Four days later, we loaded progeny from each individual into a microfluidic chip for imaging as we describe below. We imaged animals that were detectably heterozygous for reporter alleles and manually compared patterns of expression to parents. There is an obvious lack of correlation between parents and progeny, shown in Figure 8, which is representative of the individuals we imaged over three independent experiments.

### Reciprocal Crosses/maternal or paternal germline contribution

To determine if the H3K9 HMT catalytic SET domain of MET-2 from maternal and paternal germ cells was acting to regulate monoallelic expression in the early embryo, specifically the E-cell, we conducted crosses with *met-2(wam406);P_hsp-90_::mEGFP* homozygous males with *P_hsp-90_::mCherry* homozygous hermaphrodites. We performed reciprocal crosses with *P_hsp-90_::mEGFP* homozygous males crossed to *met-2(wam406);P_hsp-90_::mCherry* homozygous hermaphrodites.

To determine the contribution of set-25cat via maternal and paternal germline, we conducted crosses with *met-2(wam406);P_hsp-90_::mEGFP* homozygous males with *met-2(wam406);set-25(wam404);P_hsp-90_::mCherry* homozygous hermaphrodites. We performed reciprocal crosses with *met-2(wam406);set-25(wam404);P_hsp-90_::mEGFP* homozygous males crossed to *met-2(wam406);P_hsp-90_::mCherry* homozygous hermaphrodites.

### Microscopy

We washed day 2 adult animals (second day of adulthood at 20°) into S-basal media with tricaine/tetramisole^35^, and loaded animals into 80-lane microfluidic devices^76^. These devices immobilize worms in 80 separate lanes in a relatively restricted position, making presentation of the animals to the objective more uniform than using traditional agarose based slides. We imaged only those animals that randomly immobilized with their left side facing the cover slip, to which the fluidic device was bonded, which put intestine cells in rings I through IV closest to the microscope objective. Doing this avoids quantification error due to loss of signal with depth of tissue (i.e., imaging intestine through the germline when animals orient on their right sides).

We imaged animals as we previously described^35^. Briefly, we used a 40X 1.2 NA water objective on a Zeiss LSM780 confocal microscope. We excited the sample with 488 and 561 nm lasers and collected light from 490-550nm for mEGFP signal, and from 580-640nm for mCherry signal. For mTagBFP2 and mNeptune, we excited samples with 405 and 561nm lasers and collected emission from 410-480nm and 570-650nm, respectively. We also collected transmitted light signal for Nomarski DIC images to aid in cell identification as needed. We focused on the same field of view for each animal-starting from the posterior of the pharynx to the first half of cells in intestinal ring IV. We collected images of the entire z depth of each animal, from one side to the other, using two micrometer step size and a two micrometer optical slice as we have previously described^35^. Imaging settings were held constant between control and experimental groups.

### Image Cytometry

We conducted image cytometry in a manner that we have previously described in detail^35^. Briefly, our image cytometry consists of manual cell identification and annotation, with a semiautomatic quantification step. We first determined the orientation of the animals in images and then identified individual intestine or muscle cells. We then measured signal within an equatorial slice of the cell’s nucleus, as a proxy for the whole cell. Nuclear signal of freely diffusing monomeric fluorescent protein is nearly perfectly correlated with the cytoplasmic contents^35^. We used ImageJ software as well as custom built Nuclear Quantification Support Plugin called C. Entmoot (Alexander Seewald, Seewald Solutions, Inc., Vienna, Austria) for nucleus segmentation and signal quantification^36^.

### Data Processing and Noise Calculations

Here we measured intrinsic noise by measuring the expression level of differently colored reporter alleles in two-day old adult animals that are in a steady-state of gene expression^36^. Intrinsic noise is essentially the quantitative measure of relative deviation from the 1:1 ratio; data points having a 1:1 ratio fall on a 45° diagonal trend line. Intrinsic noise measures how deviant a pair of reporter alleles is from the average ratio among groups of cells, thus quantifying how probable it is to observe biased or monoallelic expression for a given gene (pair of alleles) in a given population of cells (e.g., muscle cells or intestine cells). The assumptions of our intrinsic noise model are the same as the assumptions in^35^. We sometimes used 8-bit or 16-bit file settings during data collection, though this difference was obviated after normalization. We normalized expression level data for each allele to per-experiment means as in previous investigations^77–79^. We calculated intrinsic noise as detailed in^77–79^. Specifically, the formula for calculating intrinsic noise is:

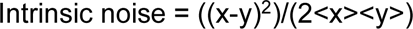

where x and y are each cell’s allele expression values and <X> and <Y> are the average value for each allele. X,Y expression data and calculated noise values for each figure are available in Source Data File 1, in Excel format. Numbers of cells and experiments are detailed in Table 2.

### Plotting and Statistics

We used SigmaPlot 12.5 (Systat Software, Inc., San Jose) for all plotting and statistical analyses. All data was non-normally distributed, even after attempting log or natural log transformations, thereby requiring nonparametric statistics for analysis. For experiments with multiple groups analyzing alleles in intestine cells or different promoters in intestine cells, we ran ANOVA on Ranks followed by Dunn’s pairwise comparisons, unless otherwise noted. For all other experiments with only two groups, we ran a nonparametric Mann Whitney U test for each distinct set of experiments. Details of each test are shown in Supplementary Information.

## Data Availability

Source data are provided with this paper. Additional data are found in main text and in Source Data. All relevant data are available from the corresponding author upon reasonable request.

## Author contributions

AM and BS designed the study. BS, AM and SY performed experiments. BS and AM analyzed the data. BS and AM wrote the initial manuscript. BS, SY, JO and AM revised the manuscript. AM and JO provided funding through grants.

## Materials availability

Material requests should be addressed to AM.

