## Supporting Information for "Maternal histone methyltransferases antagonistically regulate monoallelic expression in *C. elegans*"

### For

### Sands et al, 2024

**Supplementary Information Table of Contents**

**Section 1. Figures and Tables**……………………………………………………...2-12

Figure 1. Reporter Alleles, Expression Patterns and Quantification…...........................2

Figure 2. Quantification of MAE in *nrde-3(RNAi)*……………………………………….3

Figure 3. *met-2(wam007)* null suppressor screen…………………………………….…..4

Figure 4. Quantification of MAE in *met-2(wam007);hpl-2(RNAi)*…………………………..5

Figure 5. Lifespan Boxplots....………..………………………………………………….6

Table 1. Strains………………………………………………………………………….7

Table 2. CRISPR details ……………………………………………………………….9

Table 3. List of primers………………………………………………………………..10

Table 4. Crosses……………………………………………………………………….12

**Section 2. Statistics**……………………………………………………………...13-24

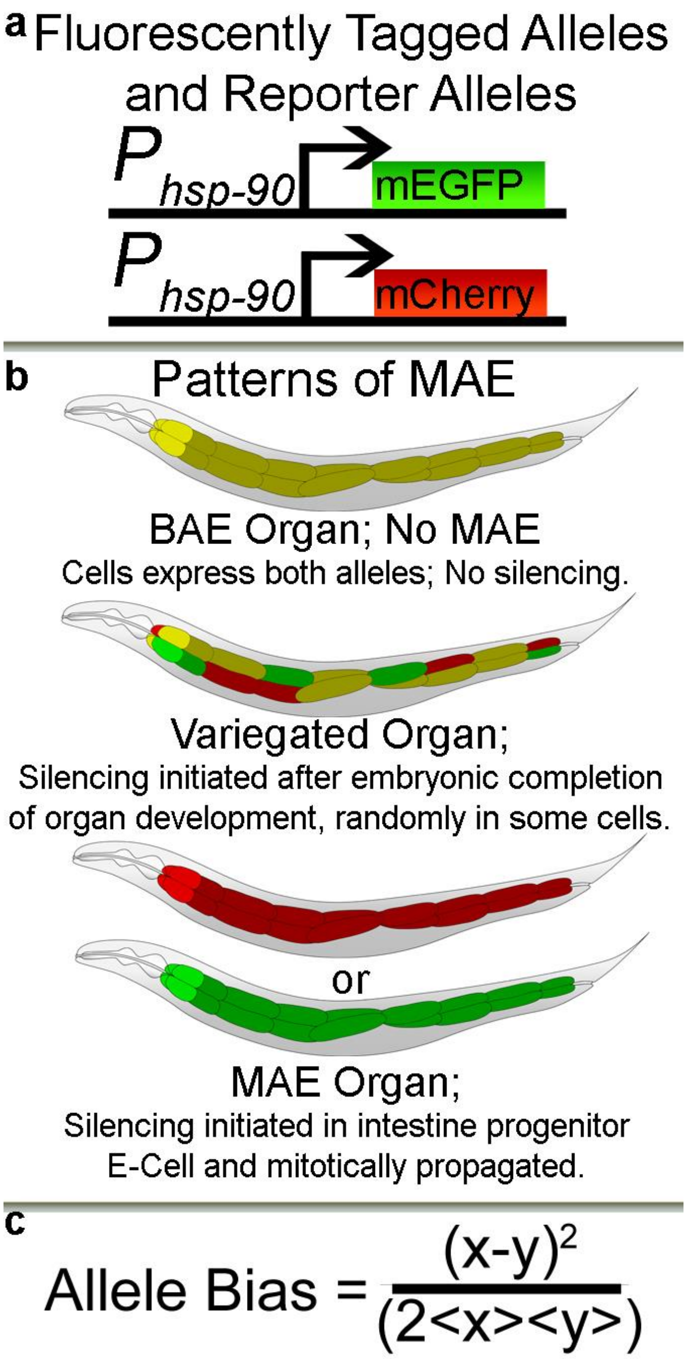


**Supplemental Figure 1. Animal Model for *In Vivo* Study of MAE, related to Figure 1. a)** We tag native alleles and generate reporter alleles. Schematic shows *hsp-90* promoter controlling expression of fluorescent reporter alleles. **b)** We quantify allele expression levels *in vivo* in the 20 intestine cells of live *C. elegans* on a point scanning confocal microscope. Cartoons show biallelic expression and different kinds of monoallelic expression. **c)** We quantify the degree of allele expression bias using the intrinsic noise formula developed by Elowitz and Swain in 2002.

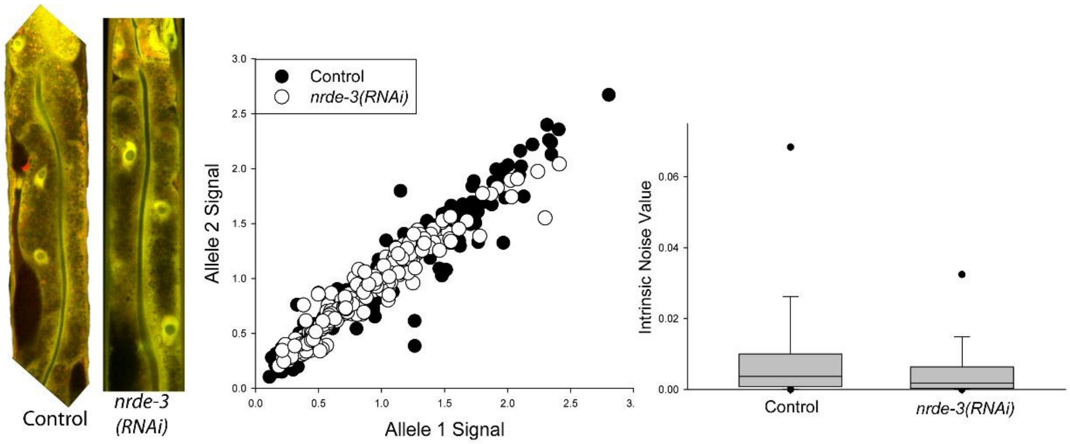


**Supplemental Figure 2. Effect of nrde-3(RNAi) on MAE.** Left: Images show $P_{hsp\text{-}90}$ reporter allele strain on EV or *nrde-3(RNAi).* Middle: Scatter plot of all cells, at least 70 cells per replicate, 3 replicates per condition. Right: Box plots of noise calculations show we detected no significant different in intrinsic noise between EV and *nrde-3(RNAi)*. While *nrde-3(RNAi)* effects approached significance, the *nrde-3(RNAi)* did not suppress the enhanced MAE in *met-2(wam007)* animals; see Supplemental Figure 3.

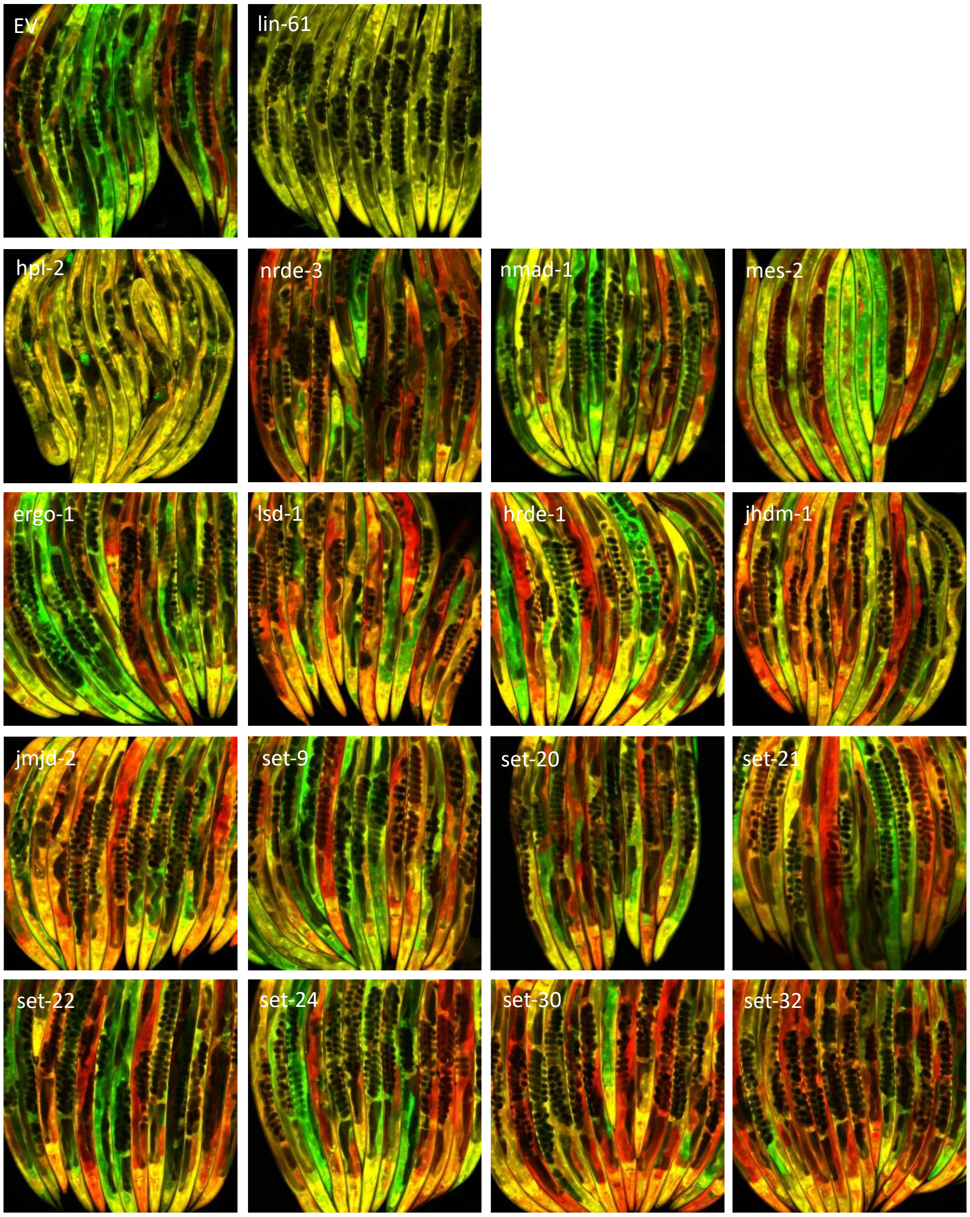


**Supplemental Figure 3**. **Suppressor screen in *met-2(wam007)* mutants**. Images are from RNAi screen of *met-2(wam007)* animals on indicated RNAi food. RNAi was conducted as described in Methods section. Images are from a Zeiss LSM 780 confocal microscope with a 10x air objective. Worms were anesthetized in round bottom 96 well plates, then transferred to a cover slip for imaging. Empty vector and *lin-61(RNAi)* were used as controls. Empty vector has no effect on MAE. *Lin-61* is a suppressor of MAE in the met-2 null background, indicated by completely yellow worms expressing equal amounts of GFP and mCherry. The genes screened here all showed extreme BAE in our initial screen, similar to *set-25*.

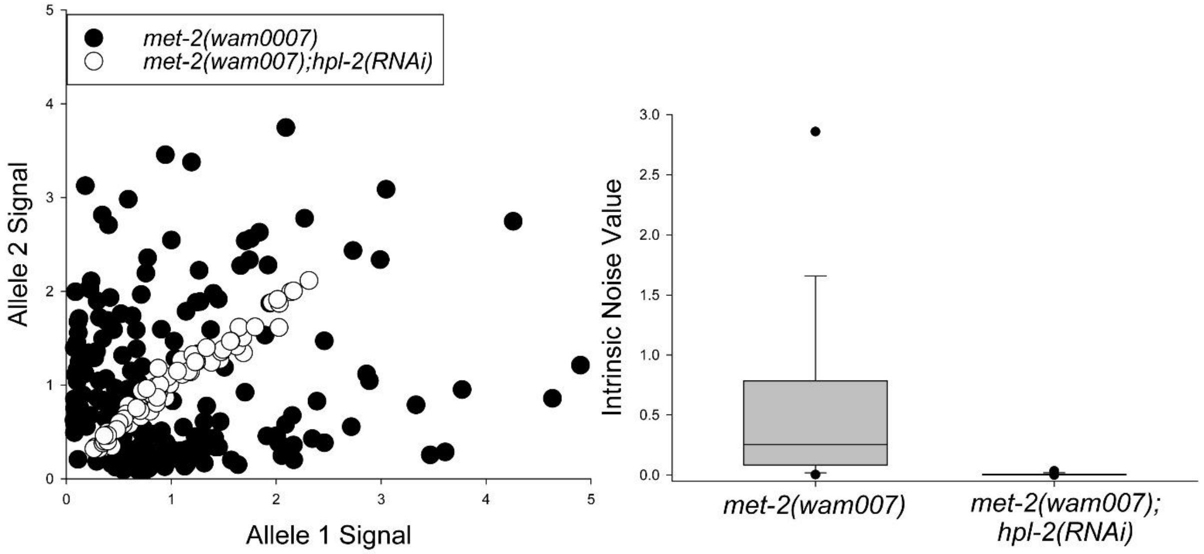


**Supplemental Figure 4. Effects of *hpl-2* on *hsp-90* reporter allele expression in *met-2(wam007)* mutants.** Left panel shows a scatter plot of *met-2(wam007)* animals' intestine cells plotted by allele expression level, compared to intestine cells in *met-2(wam007);hpl-2(RNAi)* animals. Right panel shows intrinsic noise quantified from the cells in the right panel. There was a significant difference in intrinsic noise. $P < 0.05$, ANOVA on Ranks followed by Dunn's method for multiple comparison; at least 70 cells per group measured from at least 10 individual animals. *met-2(wam007)* worms on *hpl-2(RNAi)* were imaged at 10x showing biallelic expression, similar to *set-25* and *lin-61* RNAi (see Supplemental Figure 3 and Figure 4). Here, we quantified expression from intestine as described in main text.

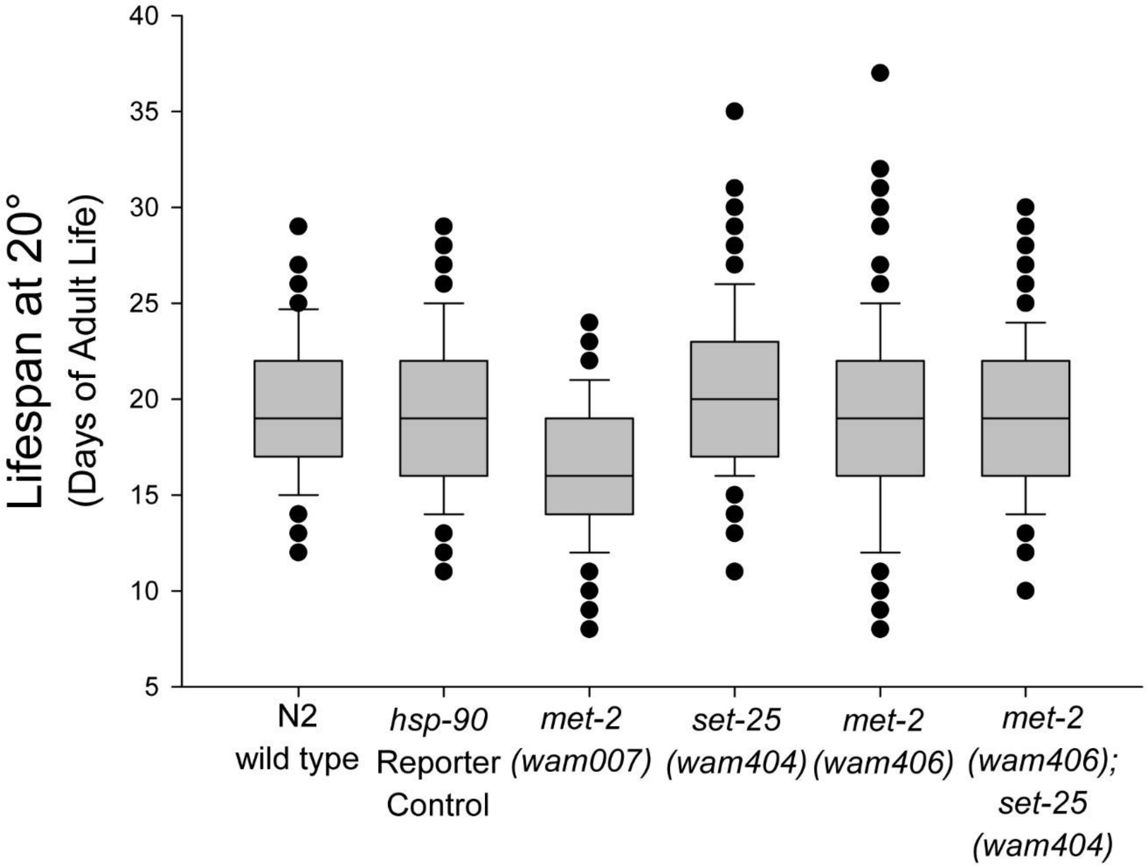


**Supplemental Figure 5. Boxplots of lifespans from Figure 6, related to Figure 6.** Boxplots show median lifespan as a line in the box, 25th and 75th percentile as vertical box bounds, 10th and 90th percentile as whisker bounds, and outliers as dots. See Analyses for all Figures section, Figure 6G for N per group.

**Supplemental Table 1. Strains**

| Strain Name | Genotype | Reference |
|---|---|---|
| ARM133 | *hutSi2661[unc-119(+), $P_{hsp-90}$:: megfp w/ 3 synthetic introns::$T_{unc-54}$, II:8420158]* | Sands et al., 2021 |
| ARM135 | *hutSi2642[unc-119(+), $P_{hsp-90}$::mcherry w/3 synthetic introns::$T_{unc-54}$, II:8420158]* | Sands et al., 2021 |
| ARM148 | *hutSi2581[unc-119(+), $P_{vit-2}$::mcherry w/ 3 synthetic introns::$T_{unc-54}$, II:8420158]* | Sands et al., 2021 |
| ARM146 | *hutSi2621[unc-119(+), $P_{vit-2}$::megfp w/ 3 synthetic introns::$T_{unc-54}$, II:8420158]* | Sands et al., 2021 |
| ARM140 | *hutSi2561[unc-119(+), $P_{hsp-16.2}$::mcherry w/ 3 synthetic introns::$T_{unc-54}$, II:8420158]* | Sands et al., 2021 |
| ARM141 | *hutSi2601[unc-119(+), $P_{hsp-16.2}$::megfp w/ 3 synthetic introns::$T_{unc-54}$, II:8420158]* | Sands et al., 2021 |
| ARM284 | *wamSi284[unc-119(+), $P_{hsp-90}$::mcherry w/3 synthetic introns::$T_{unc-54,}$ V:8643273]* | Sands et al., 2021 |
| ARM291 | *wamSi291[unc-119(+), $P_{hsp-90}$::megfp w/3 synthetic introns::$T_{unc-54,}$ V:8643273]* | Sands et al., 2021 |
| ARM268 | *wamSi268[unc-119(+), $P_{hsp-90}$::hsp-90 natural introns:: t2a :: mcherry ::$T_{unc-54,}$ II:8420158]* | Sands et al., 2021 |
| ARM263 | *wamSi263[unc-119(+), $P_{hsp-90}$::hsp-90 natural introns::t2a:: megfp ::$T_{unc-54,}$ II:8420158]* | Sands et al., 2021 |
| ARM6 | *wamSi6[unc-119(+), $P_{eef-1A.1}$::mtagBFP2 w/ 3 synthetic introns::$T_{unc-54}$, II:8420158]* | Sands et al., 2018 |
| ARM3 | *wamSi3[unc-119(+), $P_{eef-1A.1}$::mNeptune w/ 3 synthetic introns::$T_{unc-54}$, II:8420158]* | Sands et al., 2018 |
| ARM366 | *Idh-1(wam366[idh-1::T2A::mEGFP])* | This work |
| ARM379 | *Idh-1(wam379[idh-1::T2A::mCherry])* | This work |
| ARM243 | *met-2(wam007[I15stop]; [S17stop])* | This work |

| | | |
|---|---|---|
| ARM406 | *met-2(wam406[C1237A[)* | This work |
| ARM404 | *set-25(wam404[C645A])* | This work |
| ARM413 | *met-2(wam406[C1237A]; set-25(wam404[C645A])* | This work |

**Supplemental Table 2. Primers**

| Name | Sequence | Info |
|---|---|---|
| AMO328 | CGTAGATACGAGCAGTATTCTTTCG | Met-2 forward |
| AMO329 | CGTAGATACGAGCAGTTAACTTTAA | Met-2 STOP reverse |
| AMO330 | CGCCTCTTCCTGTTGCTTG | Met-2 wt reverse |
| AMO545 | GGATGATTTGGCTGATGAACTAAG | Met-2 forward |
| AMO546 | CACATGAACGTTCGGATCTGC | Met-2 cat reverse |
| AMO547 | GTGCACATTCGGATCGCA | Met-2 wt reverse |
| AMO548 | CGTTCAATGCGATGGATACTAAG | Set-25 forward |
| AMO549 | CCGAGCTGGGGTCACAG | Set-25 wt reverse |
| AMO550 | CACACTCGAAGGGTCAGC | Set-25 cat reverse |
| AMO461 | AAATTGGCTGAGAACCTCGCCAAGAAGCAAGCCCATGGATCTGGAGAGGGACGT | IDH-1 forward |
| AMO462 | GCGAGATTTTTAGATTGGGAATGAGAAAGTACTTACTTATACAATTCATCCATGCCAC | IDH-1 cherry reverse |
| AMO463 | GCGAGATTTTTAGATTGGGAATGAGAAAGTACTTATTTGTATAGTTCATCCATGCCATG | IDH-1 GFP reverse |
| AMO478 | GGGAATGAGAAAGTACTTAATGGG | IDH-1 wt reverse |

Supplemental Table 3. CRISPR details

Edit:
met-2STOP
crRNA target:
AGATACGAGCAGTATTCTTTCGG
Repair Template:
ACAGCAGTGACGAATGAACTTTGTTCCTGTGTTTCCATCCCATCATCTTAAAGTTAACTGCTCGTATC
TACGTTATTCGATGGTTCTTGTTGGTCCATCT

Edit:
met-2cat
crRNA target:
ACGTGTTGAACGTGCACATTCGG
Repair Template:
GATACAATCATAAATTTTCGATAACTTTCAGATTCTTGAATCACTCTGCAGATCCGAACGTTCATGTGC
AGCATGTCATGTACGATACGCATGATCTTCGTCTTCCATGG

Edit:
Set-25cat
crRNA target:
ACTTCGACGAACACCGAGCTGGG
Repair Template:
GCAATAAAAAATTATTTCAGGAATATCTCCCGATTCATCAATCACAGCGCTGACCCTTCGAGTGTGTTC
GTCGAAGTCTACAGTCGACGATTCGAAGAAGATCCACTGATTCCAC

Edit:
Idh-1::GFP
crRNA target:
GGAATGAGAAAGTACTTAATGGG
Repair Template:
AAATTGGCTGAGAACCTCGCCAAGAAGCAAGCCCATGGATCTGGAGAGGGACGTGGATCCCTTCTTACC
TGCGGAGACGTCGAGGAGAACCCAGGACCAATGAGTAAAGGAGAAGAACTTTTCACTGGAGTTGTCCC
AATTCTTGTTGAATTAGATGGTGATGTTAATGGGCACAAATTTTCTGTCAGTGGAGAGGGTGAAGGTGA
TGCAACATACGGAAAACTTACCCTTAAATTTATTTGCACTACTGGAAAACTACCTGTTCCATGGCCAACAC
TTGTCACTACTTTCACTTATGGTGTTCAATGCTTTTCAAGATACCCAGATCATATGAAACGGCATGACTTT
TTCAAGAGTGCCATGCCCGAAGGTTATGTACAGGAAAGAACTATATTTTTCAAAGATGACGGGAACTAC
AAGACACGTGCTGAAGTCAAGTTTGAAGGTGATACCCTTGTTAATAGAATCGAGTTAAAAGGTATTGAT
TTTAAAGAAGATGGAAACATTCTTGGACACAAATTGGAATACAACTATAACTCACACAATGTATACATCA
TGGCAGACAAACAAAAGAATGGAATCAAAGTTAACTTCAAAACTAGACACAACATTGAAGATGGAAGC
GTTCAACTAGCAGACCATTATCAACAAAATACTCCAATTGGCGATGGCCCTGTCCTTTTACCAGACAACC
ATTACCTGTCCACACAATCTAAGCTTTCGAAAGATCCCAACGAAAAGAGAGACCACATGGTCCTTCTTGA
GTTTGTAACAGCTGCTGGGATTACACATGGCATGGATGAACTATACAAATAAGTACTTTCTCATTCCCAA
TCTAAAAATCTCGC

Edit:
Idh-1::cherry
crRNA target:
GGAATGAGAAAGTACTTAAT<u>GGG</u>
Repair Template:
AAATTGGCTGAGAACCTCGCCAAGAAGCAAGCCCATGGATCTGGAGAGGGACGTGGATCCCTTCTTACC
TGCGGAGACGTCGAGGAGAACCCAGGACCAATGGTCTCAAAGGGTGAAGAAGATAACATGGCAATTAT
TAAAGAGTTTATGCGTTTCAAGGTGCATATGGAGGGATCTGTCAATGGGCATGAGTTTGAAATTGAAGG
TGAAGGAGAAGGCCGACCATATGAGGGAACACAAACCGCAAAACTAAAGGTAACTAAAGGCGGACCA
TTACCATTCGCCTGGGACATCCTCTCTCCACAGTTCATGTATGGAAGTAAAGCTTATGTTAAACATCCGGC
AGATATACCAGATTATTTGAAACTTTCATTCCCGGAGGGTTTTAAGTGGGAACGCGTAATGAATTTTGAA
GACGGAGGAGTTGTTACAGTGACGCAAGACTCAAGCCTCCAAGATGGAGAATTTATTTATAAAGTCAAA
CTTCGAGGAACGAATTTCCCCTCGGATGGACCTGTTATGCAGAAGAAGACTATGGGATGGGAAGCTTCA
AGTGAAAGAATGTACCCTGAAGACGGTGCTCTTAAGGGAGAGATTAAACAACGTCTTAAATTGAAAGAT
GGAGGACATTACGATGCTGAGGTGAAGACAACTTACAAAGCCAAAAAACCAGTTCAGCTGCCAGGAGC
GTACAATGTTAATATTAAACTGGATATCACCTCCCACAACGAGGATTACACTATCGTTGAGCAATATGAA
AGAGCTGAAGGGCGGCACTCGACAGGTGGCATGGATGAATTGTATAAGTAAGTACTTTCTCAT

Supplemental Table 4. Crosses

| Crosses | Description of heterozygous animals |
|---|---|
| ARM135 x ARM133♂ | *hsp-90* @ chr. II locus |
| ARM148 x ARM146♂ | *vit-2* @ chr. II locus |
| ARM140 x ARM141♂ | hsp-16.2 @ chr. II locus |
| ARM284 x ARM291♂ | hsp-90 @ chr. V locus |
| ARM268 x ARM263♂ | HSP90-T2A @ chr. II locus |
| ARM3 x ARM6♂ | eef-1A.1 @ chr. II locus |
| ARM406♂ x N2 | Paternal met-2 contribution |
| ARM406 x N2♂ | Maternal met-2 contribution |
| ARM406♂ x ARM413 | Maternal set-25 contribution |
| ARM406 x ARM413♂ | Paternal set-25 contribution |

#### Section 2 Statistical Analyses for All Figures

##### Figures 1 and 4 and Supplemental Figure 2

**One Way Analysis of Variance Data source:** EV RNAi screeen noise in Master stats.JNB

**Normality Test (Shapiro-Wilk)** Failed (P < 0.050)

**Kruskal-Wallis One Way Analysis of Variance on Ranks** Sunday, January 14, 2024, 5:14:02 PM

**Data source:** EV RNAi screeen noise in Master stats.JNB

| Group | N | Missing | Median | 25% | 75% |
|---|---|---|---|---|---|
| EV | 207 | 0 | 0.00371 | 0.000859 | 0.0101 |
| met-2 | 208 | 0 | 0.0607 | 0.00748 | 0.253 |
| set-25 | 208 | 0 | 0.00109 | 0.000311 | 0.00299 |
| lin-65 | 207 | 0 | 0.122 | 0.0257 | 0.497 |
| arle-14 | 210 | 0 | 0.0110 | 0.00235 | 0.0250 |
| lin-61 | 210 | 0 | 0.00105 | 0.000322 | 0.00289 |
| cec-4 | 210 | 0 | 0.00467 | 0.00106 | 0.0123 |
| lem-2 | 210 | 0 | 0.00624 | 0.00125 | 0.0209 |
| nrde-3 | 209 | 0 | 0.00189 | 0.000335 | 0.00639 |

H = 611.498 with 8 degrees of freedom. (P = <0.001)

The differences in the median values among the treatment groups are greater than would be expected by chance; there is a statistically significant difference (P = <0.001)

To isolate the group or groups that differ from the others use a multiple comparison procedure.

All Pairwise Multiple Comparison Procedures (Dunn's Method) :

| Comparison | Diff of Ranks | Q | P<0.05 |
|---|---|---|---|
| lin-65 vs lin-61 | 934.564 | 17.587 | Yes |
| lin-65 vs set-25 | 920.185 | 17.275 | Yes |
| lin-65 vs nrde-3 | 815.497 | 15.328 | Yes |
| lin-65 vs EV | 665.715 | 12.483 | Yes |
| lin-65 vs cec-4 | 607.440 | 11.431 | Yes |
| lin-65 vs lem-2 | 530.421 | 9.982 | Yes |
| lin-65 vs arle-14 | 436.736 | 8.219 | Yes |
| lin-65 vs met-2 | 127.848 | 2.400 | No |
| met-2 vs lin-61 | 806.716 | 15.199 | Yes |
| met-2 vs set-25 | 792.337 | 14.893 | Yes |
| met-2 vs nrde-3 | 687.649 | 12.941 | Yes |
| met-2 vs EV | 537.866 | 10.098 | Yes |
| met-2 vs cec-4 | 479.592 | 9.036 | Yes |
| met-2 vs lem-2 | 402.573 | 7.585 | Yes |
| met-2 vs arle-14 | 308.887 | 5.820 | Yes |
| arle-14 vs lin-61 | 497.829 | 9.402 | Yes |
| arle-14 vs set-25 | 483.449 | 9.109 | Yes |
| arle-14 vs nrde-3 | 378.762 | 7.145 | Yes |
| arle-14 vs EV | 228.979 | 4.309 | Yes |
| arle-14 vs cec-4 | 170.705 | 3.224 | Yes |
| arle-14 vs lem-2 | 93.686 | 1.769 | No |
| lem-2 vs lin-61 | 404.143 | 7.633 | Yes |

| | | | |
|---|---|---|---|
| lem-2 vs set-25 | 389.764 | 7.343 | Yes |
| lem-2 vs nrde-3 | 285.076 | 5.378 | Yes |
| lem-2 vs EV | 135.294 | 2.546 | No |
| lem-2 vs cec-4 | 77.019 | 1.455 | Do Not Test |
| cec-4 vs lin-61 | 327.124 | 6.178 | Yes |
| cec-4 vs set-25 | 312.745 | 5.892 | Yes |
| cec-4 vs nrde-3 | 208.057 | 3.925 | Yes |
| cec-4 vs EV | 58.275 | 1.097 | Do Not Test |
| EV vs lin-61 | 268.849 | 5.059 | Yes |
| EV vs set-25 | 254.470 | 4.777 | Yes |
| EV vs nrde-3 | .714983 | 2.815 | No |
| nrde-3 vs lin-61 | 119.067 | 2.246 | No |
| nrde-3 vs set-25 | 104.688 | 1.970 | Do Not Test |
| set-25 vs lin-61 | 14.379 | 0.271 | Do Not Test |

Note: The multiple comparisons on ranks do not include an adjustment for ties.

---

Figures 4g&h and Supplemental Figure 4

**Normality Test (Shapiro-Wilk)** Failed (P < 0.050)

| **Group** | **N** | **Missing** | **Median** | **25%** | **75%** |
|---|---|---|---|---|---|
| met-2ko | 199 | 0 | 0.254 | 0.0840 | 0.784 |
| met-2ko;set-25RNAi | 204 | 0 | 0.00237 | 0.000525 | 0.0819 |
| met2 lin61 | 210 | 0 | 0.00160 | 0.000328 | 0.00412 |
| met-2; hpl-2rnai | 70 | 0 | 0.00363 | 0.000912 | 0.00854 |

H = 285.132 with 3 degrees of freedom. (P = <0.001)

The differences in the median values among the treatment groups are greater than would be expected by chance; there is a statistically significant difference (P = <0.001)

To isolate the group or groups that differ from the others use a multiple comparison procedure.

All Pairwise Multiple Comparison Procedures (Dunn's Method) :

| **Comparison** | **Diff of Ranks** | **Q** | **P<0.05** |
|---|---|---|---|
| met-2ko vs met2 lin61 | 314.943 | 16.135 | Yes |
| met-2ko vs met-2; hpl-2rnai | 258.924 | 9.443 | Yes |
| met-2ko vs met-2ko;set-25RNAi | 228.846 | 11.641 | Yes |
| met-2ko;set-2 vs met2 lin61 | 86.096 | 4.439 | Yes |
| met-2ko;set-2 vs met-2; hpl-2r | 30.077 | 1.100 | No |
| met-2; hpl-2rnai vs met2 lin61 | 56.019 | 2.057 | No |

Note: The multiple comparisons on ranks do not include an adjustment for ties.

-----------------------------------------------------------------------------------------------------------------------------------------

---

Figure 2a

**Mann-Whitney Rank Sum Test**

**Normality Test (Shapiro-Wilk)** Failed (P < 0.050)

| Group | N | Missing | Median | 25% | 75% |
|---|---|---|---|---|---|
| Control | 209 | 0 | 0.00232 | 0.000737 | 0.00770 |
| met-2 ko | 208 | 0 | 0.187 | 0.0318 | 0.465 |

Mann-Whitney U Statistic= 4353.000

T = 60855.000 n(small)= 208 n(big)= 209 (P = <0.001)

The difference in the median values between the two groups is greater than would be expected by chance; there is a statistically significant difference (P = <0.001)

---

Figure 2c

**Mann-Whitney Rank Sum Test**

**Normality Test (Shapiro-Wilk)** Failed (P < 0.050)

| Group | N | Missing | Median | 25% | 75% |
|---|---|---|---|---|---|
| control t2a | 196 | 0 | 0.00163 | 0.000292 | 0.00611 |
| met-2 ko t2a | 210 | 0 | 0.100 | 0.0166 | 0.352 |

Mann-Whitney U Statistic= 4215.000

T = 23521.000 n(small)= 196 n(big)= 210 (P = <0.001)

The difference in the median values between the two groups is greater than would be expected by chance; there is a statistically significant difference (P = <0.001)

---

Figure 2b

**Mann-Whitney Rank Sum Test**

**Normality Test (Shapiro-Wilk)** Failed (P < 0.050)

| Group | N | Missing | Median | 25% | 75% |
|---|---|---|---|---|---|
| EV chr V | 207 | 0 | 0.0196 | 0.00363 | 0.0519 |
| CHR V met-2 RNAi | 206 | 0 | 0.180 | 0.0552 | 0.551 |

Mann-Whitney U Statistic= 8475.000

T = 55488.000 n(small)= 206 n(big)= 207 (P = <0.001)

The difference in the median values between the two groups is greater than would be expected by chance; there is a statistically significant difference (P = <0.001)

---

Figure 3a

**Mann-Whitney Rank Sum Test**

**Normality Test (Shapiro-Wilk)** Failed (P < 0.050)

| Group | N | Missing | Median | 25% | 75% |
|---|---|---|---|---|---|
| vit-2 | 210 | 0 | 0.000968 | 0.000319 | 0.00436 |
| met-2 ko vit2 | 209 | 0 | 0.000809 | 0.000246 | 0.00275 |

Mann-Whitney U Statistic= 20567.000

T = 42512.000  n(small)= 209  n(big)= 210  (P = 0.266)

The difference in the median values between the two groups is not great enough to exclude the possibility that the difference is due to random sampling variability; there is not a statistically significant difference (P = 0.266)

---

Figure 3b

**Mann-Whitney Rank Sum Test**

**Data source:** met-2 ko chrV n other genes noise in Master stats.JNB

**Normality Test (Shapiro-Wilk)** Failed  (P < 0.050)

| **Group** | **N** | **Missing** | **Median** | **25%** | **75%** |
|---|---|---|---|---|---|
| 16.2 ev | 210 | 0 | 0.00297 | 0.000699 | 0.00783 |
| 16.2 met2 RNAi | 210 | 0 | 0.00311 | 0.000754 | 0.00994 |

Mann-Whitney U Statistic= 21389.000

T = 43544.000  n(small)= 210  n(big)= 210  (P = 0.595)

The difference in the median values between the two groups is not great enough to exclude the possibility that the difference is due to random sampling variability; there is not a statistically significant difference (P = 0.595)

---

Figure 3c

**Mann-Whitney Rank Sum Test**

**Normality Test (Shapiro-Wilk)** Failed  (P < 0.050)

| **Group** | **N** | **Missing** | **Median** | **25%** | **75%** |
|---|---|---|---|---|---|
| idh-1 ev | 208 | 0 | 0.00356 | 0.00104 | 0.0123 |
| idh-1 met-2 RNAi | 209 | 0 | 0.00337 | 0.000888 | 0.0151 |

Mann-Whitney U Statistic= 21311.000

T = 43897.000  n(small)= 208  n(big)= 209  (P = 0.730)

The difference in the median values between the two groups is not great enough to exclude the possibility that the difference is due to random sampling variability; there is not a statistically significant difference (P = 0.730)

---

Figure 3d

**Mann-Whitney Rank Sum Test**

**Data source:** met-2 ko chrV n other genes noise in Master stats.JNB

**Normality Test (Shapiro-Wilk)** Failed (P < 0.050)

| Group | N | Missing | Median | 25% | 75% |
|---|---|---|---|---|---|
| EV EFT-3 | 180 | 0 | 0.00254 | 0.000697 | 0.00730 |
| eft-3 met-2 RNAi | 180 | 0 | 0.00944 | 0.00248 | 0.0339 |

Mann-Whitney U Statistic= 9359.000

T = 25649.000 n(small)= 180 n(big)= 180 (P = <0.001)

The difference in the median values between the two groups is greater than would be expected by chance; there is a statistically significant difference (P = <0.001)

---

Figure 5

**Kruskal-Wallis One Way Analysis of Variance on Ranks**
**Normality Test (Shapiro-Wilk)** Failed (P < 0.050)

| Group | N | Missing | Median | 25% | 75% |
|---|---|---|---|---|---|
| control | 209 | 0 | 0.00459 | 0.000925 | 0.0144 |
| 404 | 210 | 0 | 0.00114 | 0.000288 | 0.00381 |
| 406 | 210 | 0 | 0.134 | 0.0237 | 0.469 |
| 414 | 210 | 0 | 0.00140 | 0.000351 | 0.00438 |

H = 355.298 with 3 degrees of freedom. (P = <0.001)

The differences in the median values among the treatment groups are greater than would be expected by chance; there is a statistically significant difference (P = <0.001)

To isolate the group or groups that differ from the others use a multiple comparison procedure.

All Pairwise Multiple Comparison Procedures (Dunn's Method) :

| Comparison | Diff of Ranks | Q | P<0.05 |
|---|---|---|---|
| 406 vs 404 | 396.676 | 16.773 | Yes |
| 406 vs 414 | 374.019 | 15.815 | Yes |
| 406 vs control | 269.196 | 11.369 | Yes |
| control vs 404 | 127.480 | 5.384 | Yes |
| control vs 414 | 104.823 | 4.427 | Yes |
| 414 vs 404 | 22.657 | 0.958 | No |

Note: The multiple comparisons on ranks do not include an adjustment for ties.

---

Figure 6a

Day 1 Adults

**Kruskal-Wallis One Way Analysis of Variance on Ranks**
**Data source:** daily progeny production v 2 in Master stats.JNB

**Normality Test (Shapiro-Wilk)** Passed (P = 0.572)

**Equal Variance Test:** Failed (P < 0.050)

| Group | N | Missing | Median | 25% | 75% |
|---|---|---|---|---|---|
| hsp90 | 18 | 0 | 100.500 | 94.000 | 113.500 |
| met-2KO | 18 | 0 | 2.000 | 0.000 | 15.500 |
| met-2cat | 18 | 0 | 81.500 | 23.750 | 96.250 |
| set-25cat | 18 | 0 | 106.500 | 90.750 | 115.750 |
| doublecat | 18 | 0 | 111.500 | 92.500 | 124.500 |

H = 53.867 with 4 degrees of freedom. (P = <0.001)

The differences in the median values among the treatment groups are greater than would be expected by chance; there is a statistically significant difference (P = <0.001)

To isolate the group or groups that differ from the others use a multiple comparison procedure.

All Pairwise Multiple Comparison Procedures (Tukey Test):

| Comparison | Diff of Ranks | q | P<0.05 |
|---|---|---|---|
| doublecat vs met-2KO | 946.000 | 8.535 | Yes |
| doublecat vs met-2cat | 530.000 | 4.782 | Yes |
| doublecat vs hsp90 | 132.000 | 1.191 | No |
| doublecat vs set-25cat | 24.500 | 0.221 | Do Not Test |
| set-25cat vs met-2KO | 921.500 | 8.314 | Yes |
| set-25cat vs met-2cat | 505.500 | 4.561 | Yes |
| set-25cat vs hsp90 | 107.500 | 0.970 | Do Not Test |
| hsp90 vs met-2KO | 814.000 | 7.344 | Yes |
| hsp90 vs met-2cat | 398.000 | 3.591 | No |
| met-2cat vs met-2KO | 416.000 | 3.753 | No |

Note: The multiple comparisons on ranks do not include an adjustment for ties.

A result of "Do Not Test" occurs for a comparison when no significant difference is found between the two rank sums that enclose that comparison. For example, if you had four rank sums sorted in order, and found no significant difference between rank sums 4 vs. 2, then you would not test 4 vs. 3 and 3 vs. 2, but still test 4 vs. 1 and 3 vs. 1 (4 vs. 3 and 3 vs. 2 are enclosed by 4 vs. 2: 4 3 2 1). Note that not testing the enclosed rank sums is a procedural rule, and a result of Do Not Test should be treated as if there is no significant difference between the rank sums, even though one may appear to exist.

---

Figure 6b

#### Day 2 Adults

**Kruskal-Wallis One Way Analysis of Variance on Ranks**

**Data source:** daily progeny production v 2 in Master stats.JNB

**Normality Test (Shapiro-Wilk)** Passed (P = 0.684)

**Equal Variance Test:** Failed (P < 0.050)

| Group | N | Missing | Median | 25% | 75% |
|---|---|---|---|---|---|
| hsp90 | 18 | 0 | 123.000 | 109.250 | 135.500 |
| met-2KO | 18 | 0 | 12.500 | 0.000 | 56.500 |
| met-2cat | 18 | 0 | 94.000 | 57.750 | 133.500 |
| set-25cat | 18 | 0 | 133.500 | 123.000 | 149.000 |

| | | | | | |
|---|---|---|---|---|---|
| doublecat | 18 | 0 | 104.500 | 93.250 | 125.000 |

H = 48.032 with 4 degrees of freedom. (P = <0.001)

The differences in the median values among the treatment groups are greater than would be expected by chance; there is a statistically significant difference (P = <0.001)

To isolate the group or groups that differ from the others use a multiple comparison procedure.

All Pairwise Multiple Comparison Procedures (Tukey Test):

| **Comparison** | **Diff of Ranks** | **q** | **P<0.05** |
|---|---|---|---|
| set-25cat vs met-2KO | 1023.000 | 9.230 | Yes |
| set-25cat vs met-2cat | 448.500 | 4.046 | Yes |
| set-25cat vs doublecat | 417.000 | 3.762 | No |
| set-25cat vs hsp90 | 199.000 | 1.795 | Do Not Test |
| hsp90 vs met-2KO | 824.000 | 7.434 | Yes |
| hsp90 vs met-2cat | 249.500 | 2.251 | No |
| hsp90 vs doublecat | 218.000 | 1.967 | Do Not Test |
| doublecat vs met-2KO | 606.000 | 5.467 | Yes |
| doublecat vs met-2cat | 31.500 | 0.284 | Do Not Test |
| met-2cat vs met-2KO | 574.500 | 5.183 | Yes |

Note: The multiple comparisons on ranks do not include an adjustment for ties.

A result of "Do Not Test" occurs for a comparison when no significant difference is found between the two rank sums that enclose that comparison. For example, if you had four rank sums sorted in order, and found no significant difference between rank sums 4 vs. 2, then you would not test 4 vs. 3 and 3 vs. 2, but still test 4 vs. 1 and 3 vs. 1 (4 vs. 3 and 3 vs. 2 are enclosed by 4 vs. 2: 4 3 2 1). Note that not testing the enclosed rank sums is a procedural rule, and a result of Do Not Test should be treated as if there is no significant difference between the rank sums, even though one may appear to exist.

---

Figure 6c

#### Day 3 Adults

**Kruskal-Wallis One Way Analysis of Variance on Ranks**
**Data source:** daily progeny production v 2 in Master stats.JNB

**Normality Test (Shapiro-Wilk)** Failed (P < 0.050)

| **Group** | **N** | **Missing** | **Median** | **25%** | **75%** |
|---|---|---|---|---|---|
| hsp90 | 18 | 0 | 71.500 | 43.000 | 92.500 |
| met-2KO | 18 | 0 | 9.000 | 0.000 | 45.250 |
| met-2cat | 18 | 0 | 42.500 | 13.000 | 99.750 |
| set-25cat | 18 | 0 | 64.500 | 53.500 | 81.250 |
| doublecat | 18 | 0 | 14.500 | 8.750 | 32.000 |

H = 31.125 with 4 degrees of freedom. (P = <0.001)

The differences in the median values among the treatment groups are greater than would be expected by chance; there is a statistically significant difference (P = <0.001)

To isolate the group or groups that differ from the others use a multiple comparison procedure.

All Pairwise Multiple Comparison Procedures (Student-Newman-Keuls Method) :

| **Comparison** | **Diff of Ranks** | **q** | **P<0.05** |
|---|---|---|---|
| hsp90 vs met-2KO | 639.000 | 5.765 | Yes |
| hsp90 vs doublecat | 636.500 | 7.168 | Yes |
| hsp90 vs met-2cat | 283.000 | 4.240 | Yes |
| hsp90 vs set-25cat | 44.000 | 0.984 | No |
| set-25cat vs met-2KO | 595.000 | 6.701 | Yes |
| set-25cat vs doublecat | 592.500 | 8.877 | Yes |
| set-25cat vs met-2cat | 239.000 | 5.347 | Yes |
| met-2cat vs met-2KO | 356.000 | 5.334 | Yes |
| met-2cat vs doublecat | 353.500 | 7.908 | Yes |
| doublecat vs met-2KO | 2.500 | 0.0559 | No |

Note: The multiple comparisons on ranks do not include an adjustment for ties.

---

Figure 6d

#### Day 4 Adults

**Kruskal-Wallis One Way Analysis of Variance on Ranks**
**Data source:** daily progeny production v 2 in Master stats.JNB

**Normality Test (Shapiro-Wilk)** Failed (P < 0.050)

| **Group** | **N** | **Missing** | **Median** | **25%** | **75%** |
|---|---|---|---|---|---|
| hsp90 | 18 | 0 | 6.000 | 1.750 | 11.500 |
| met-2KO | 18 | 0 | 1.000 | 0.000 | 5.500 |
| met-2cat | 18 | 0 | 2.500 | 0.000 | 16.000 |
| set-25cat | 18 | 0 | 10.000 | 5.000 | 17.000 |
| doublecat | 18 | 0 | 3.500 | 1.000 | 7.000 |

H = 12.823 with 4 degrees of freedom. (P = 0.012)

The differences in the median values among the treatment groups are greater than would be expected by chance; there is a statistically significant difference (P = 0.012)

To isolate the group or groups that differ from the others use a multiple comparison procedure.

All Pairwise Multiple Comparison Procedures (Student-Newman-Keuls Method) :

| **Comparison** | **Diff of Ranks** | **q** | **P<0.05** |
|---|---|---|---|
| set-25cat vs met-2KO | 509.000 | 4.592 | Yes |
| set-25cat vs doublecat | 380.500 | 4.285 | Yes |
| set-25cat vs met-2cat | 355.000 | 5.319 | Yes |
| set-25cat vs hsp90 | 185.500 | 4.150 | Yes |
| hsp90 vs met-2KO | 323.500 | 3.643 | Yes |
| hsp90 vs doublecat | 195.000 | 2.922 | No |
| hsp90 vs met-2cat | 169.500 | 3.792 | Do Not Test |
| met-2cat vs met-2KO | 154.000 | 2.307 | No |
| met-2cat vs doublecat | 25.500 | 0.570 | Do Not Test |
| doublecat vs met-2KO | 128.500 | 2.875 | Do Not Test |

Note: The multiple comparisons on ranks do not include an adjustment for ties.

A result of "Do Not Test" occurs for a comparison when no significant difference is found between the two rank sums that enclose that comparison. For example, if you had four rank sums sorted in order, and found no significant difference between rank sums 4 vs. 2, then you would not test 4 vs. 3 and 3 vs. 2, but still test 4 vs. 1 and 3 vs. 1 (4 vs. 3 and 3 vs. 2 are enclosed by 4 vs. 2: 4 3 2 1). Note that not testing the enclosed rank sums is a procedural rule, and a result of Do Not Test should be treated as if there is no significant difference between the rank sums, even though one may appear to exist.

---

Figure 6e

#### Total Progeny Per Animal

**One Way Analysis of Variance**
**Normality Test (Shapiro-Wilk)** Passed (P = 0.073)
**Equal Variance Test:** Failed (P < 0.050)
**Kruskal-Wallis One Way Analysis of Variance on Ranks**
**Data source:** Data 1 in Notebook1

Dependent Variable:

| Group | N | Missing | Median | 25% | 75% |
|---|---|---|---|---|---|
| control | 18 | 0 | 308.500 | 275.750 | 329.000 |
| met-2 ko | 18 | 0 | 33.000 | 0.000 | 140.000 |
| met-2 cat | 18 | 0 | 230.500 | 116.000 | 320.500 |
| set-25 cat | 18 | 0 | 325.500 | 282.750 | 349.000 |
| met-2 cat;set-25 cat | 18 | 0 | 233.500 | 212.000 | 267.500 |

H = 48.790 with 4 degrees of freedom. (P = <0.001)

The differences in the median values among the treatment groups are greater than would be expected by chance; there is a statistically significant difference (P = <0.001)

To isolate the group or groups that differ from the others use a multiple comparison procedure.

All Pairwise Multiple Comparison Procedures (Student-Newman-Keuls Method) :

| Comparison | Diff of Ranks | q | P<0.05 |
|---|---|---|---|
| set-25 cat vs met-2 ko | 989.500 | 8.927 | Yes |
| set-25 cat vs met-2 cat;set | 519.000 | 5.845 | Yes |
| set-25 cat vs met-2 cat | 441.000 | 6.607 | Yes |
| set-25 cat vs control | 123.000 | 2.752 | No |
| control vs met-2 ko | 866.500 | 9.759 | Yes |
| control vs met-2 cat;set | 396.000 | 5.933 | Yes |
| control vs met-2 cat | 318.000 | 7.114 | Yes |
| met-2 cat vs met-2 ko | 548.500 | 8.218 | Yes |
| met-2 cat vs met-2 cat;set | 78.000 | 1.745 | No |
| met-2 cat;set vs met-2 ko | 470.500 | 10.526 | Yes |

Note: The multiple comparisons on ranks do not include an adjustment for ties.

---

Figure 6f

Coefficient of Variation for Total Progeny Per Individual
**Kruskal-Wallis One Way Analysis of Variance on Ranks**
**Data source:** Fecundity in Master stats.JNB

**One Way Analysis of Variance**
**Data source:** Fecundity in Master stats.JNB

| Group Name | N | Missing | Mean | Std Dev | SEM |
|---|---|---|---|---|---|
| control | 3 | 0 | 0.131 | 0.0825 | 0.0477 |
| met-2 ko | 3 | 0 | 1.199 | 0.157 | 0.0908 |
| met-2 cat | 3 | 0 | 0.554 | 0.349 | 0.202 |
| set-25 cat | 3 | 0 | 0.106 | 0.0450 | 0.0260 |
| met-2 cat;set-25 cat | 3 | 0 | 0.151 | 0.0728 | 0.0420 |

| Source of Variation | DF | SS | MS | F | P |
|---|---|---|---|---|---|
| Between Groups | 4 | 2.633 | 0.658 | 20.462 | <0.001 |
| Residual | 10 | 0.322 | 0.0322 | | |
| Total | 14 | 2.955 | | | |

The differences in the mean values among the treatment groups are greater than would be expected by chance; there is a statistically significant difference (P = <0.001).

Power of performed test with alpha = 0.050: 1.000

Multiple Comparisons versus Control Group (Holm-Sidak method):
Overall significance level = 0.05

Comparisons for factor:

| Comparison | Diff of Means | t | P | P<0.050 |
|---|---|---|---|---|
| control vs. met-2 ko | 1.067 | 7.288 | <0.001 | Yes |
| control vs. met-2 cat | 0.423 | 2.889 | 0.048 | Yes |
| control vs. set-25 cat | 0.0248 | 0.169 | 0.983 | No |
| control vs. met-2 cat;se | 0.0203 | 0.139 | 0.892 | No |

---

Figure 6g
Lifespan
**Normality Test (Shapiro-Wilk)** Failed (P < 0.050)

**Kruskal-Wallis One Way Analysis of Variance on Ranks**

| Group | N | Missing | Median | 25% | 75% |
|---|---|---|---|---|---|
| N2 | 182 | 0 | 19.000 | 17.000 | 22.000 |
| hsp90 ctrl | 222 | 0 | 19.000 | 16.000 | 22.000 |
| met-2(wam007) | 174 | 0 | 16.000 | 14.000 | 19.000 |
| set-25(wam404) | 265 | 0 | 20.000 | 17.000 | 23.000 |
| met-2(wam406) | 160 | 0 | 19.000 | 16.000 | 22.000 |
| double cat | 231 | 0 | 19.000 | 16.000 | 22.000 |

H = 104.717 with 5 degrees of freedom. (P = <0.001)

The differences in the median values among the treatment groups are greater than would be expected by chance; there is a statistically significant difference (P = <0.001)

To isolate the group or groups that differ from the others use a multiple comparison procedure.

All Pairwise Multiple Comparison Procedures (Dunn's Method) :

| Comparison | Diff of Ranks | Q | P<0.05 |
|---|---|---|---|
| set-25(wam404 vs met-2(wam007) | 352.841 | 10.147 | Yes |
| set-25(wam404 vs met-2(wam406) | 128.733 | 3.608 | Yes |
| set-25(wam404) vs hsp90 ctrl | 119.050 | 3.672 | Yes |
| set-25(wam404) vs double cat | 118.490 | 3.694 | Yes |
| set-25(wam404) vs N2 | 113.763 | 3.316 | Yes |
| N2 vs met-2(wam007) | 239.078 | 6.327 | Yes |
| N2 vs met-2(wam406) | 14.970 | 0.388 | No |
| N2 vs hsp90 ctrl | 5.287 | 0.148 | Do Not Test |
| N2 vs double cat | 4.727 | 0.134 | Do Not Test |
| double cat vs met-2(wam007) | 234.351 | 6.551 | Yes |
| double cat vs met-2(wam406) | 10.244 | 0.279 | Do Not Test |
| double cat vs hsp90 ctrl | 0.561 | 0.0167 | Do Not Test |
| hsp90 ctrl vs met-2(wam007) | 233.790 | 6.479 | Yes |
| hsp90 ctrl vs met-2(wam406) | 9.683 | 0.262 | Do Not Test |
| met-2(wam406) vs met-2(wam007) | 224.108 | 5.741 | Yes |

Note: The multiple comparisons on ranks do not include an adjustment for ties.

--------------------------------------------------------------------------------------------------------------------------------------------

Figure 9f

**Mann-Whitney Rank Sum Test**
**Data source:** Data 11 in Master stats.JNB

**Normality Test (Shapiro-Wilk)** Failed (P < 0.050)

| Group | N | Missing | Median | 25% | 75% |
|---|---|---|---|---|---|
| m2cat f | 180 | 0 | 0.0469 | 0.00781 | 0.159 |
| m2cat m | 180 | 0 | 0.00193 | 0.000572 | 0.00737 |

Mann-Whitney U Statistic= 5438.000

T = 43252.000 n(small)= 180 n(big)= 180 (P = <0.001)

The difference in the median values between the two groups is greater than would be expected by chance; there is a statistically significant difference (P = <0.001)

---

Figure 9l
**Mann-Whitney Rank Sum Test**
**Data source:** Data 11 in Master stats.JNB

**Normality Test (Shapiro-Wilk)** Failed (P < 0.050)

| Group | N | Missing | Median | 25% | 75% |
|---|---|---|---|---|---|
| dc f | 180 | 0 | 0.00109 | 0.000308 | 0.00267 |
| dc m | 180 | 0 | 0.0508 | 0.0122 | 0.154 |

Mann-Whitney U Statistic= 2279.000

T = 18569.000  n(small)= 180  n(big)= 180  (P = <0.001)

The difference in the median values between the two groups is greater than would be expected by chance; there is a statistically significant difference  (P = <0.001)
